# NAC controls nascent chain fate through tunnel sensing and chaperone action

**DOI:** 10.1101/2025.07.27.667080

**Authors:** Jae Ho Lee, Laurenz Rabl, Martin Gamerdinger, Vaishali Goyal, Katrin Michaela Khakzar, Natalia Moreira Barbosa, Juliana Abramovich, Fabian Morales-Polanco, Ann-Kathrin Köhler, Ekaterina Samatova, Marina V. Rodnina, Elke Deuerling, Judith Frydman

**Affiliations:** Department of Biology, Stanford University, Stanford, CA 94305, USA; Department of Biology, Molecular Microbiology, University of Konstanz, 78457 Konstanz, Germany; Department of Physical Biochemistry, Max Planck Institute for Multidisciplinary Sciences, Goettingen 37077, Germany; Glenn Laboratories for the Biology of Aging, Stanford University, Stanford, CA, USA; Department of Biochemistry and Cell Biology, Stony Brook University, Stony Brook, NY 11794, USA

## Abstract

The nascent polypeptide-associated complex (NAC) is a conserved ribosome-bound factor with essential yet incompletely understood roles in protein biogenesis. Here, we show that NAC is a multifaceted regulator that coordinates translation elongation, cotranslational folding, and organelle targeting through distinct interactions with nascent polypeptides both inside and outside the ribosome exit tunnel. Using NAC-selective ribosome profiling in *C. elegans*, we identify thousands of sequence-specific NAC binding events across the nascent proteome, revealing broad cotranslational engagement with hydrophobic and helical motifs in cytosolic, nuclear, ER, and mitochondrial proteins. Unexpectedly, we discover an intra-tunnel sensing mode, where NAC engages ribosomes with extremely short nascent polypeptides inside the exit tunnel in a sequence-specific manner. These early NAC interactions induce an early elongation slowdown that tunes ribosome flux and prevent ribosome collisions, linking NAC’s chaperone activity to kinetic control of translation. We propose that NAC action protects aggregation-prone intermediates by shielding amphipathic helices thus promoting cytonuclear folding and supporting mitochondrial membrane protein biogenesis and ER targeting by early recognition of signal sequences and transmembrane domain. Our findings establish NAC as an early-acting, multifaceted orchestrator of cotranslational proteostasis, with distinct mechanisms of action on nascent chains depending on their sequence features and subcellular destinations.

## Main

Correct biogenesis of newly synthesized proteins is essential for maintaining a healthy cellular proteome^16,17^. Nascent polypeptides on translating ribosomes remain incomplete and unable to reach their stably folded final structures and are thus susceptible to misfolding and aggregation^11,18^. Translation speed is often fine-tuned to regulate the timing of different cotranslational processes^13,19^. However, cotranslational folding and transport to different organelles relies on a network of cotranslationally acting chaperones and targeting factors^12,13^. One of the early chaperones, the nascent polypeptide-associated complex (NAC) is an abundant heterodimer of NACα and NACβ^12^, essential in higher eukaryotes and implicated in various cotranslational events^20,21^. NAC interacts with translating ribosomes with nanomolar affinity^22,23^ and enhances the specificity of cotranslational targeting by signal recognition particle (SRP) to the endoplasmic reticulum (ER)^24–27^. Recent work showed NAC acts to coordinate N-terminal processing of nascent proteins by recruiting N-terminal processing methionine aminopeptidase 1 (METAP1)^28^, N-acetyltransferase A (NatA)^29^, and N-myristoyltransferases (NMTs)^30^. While general substrates of NAC have been surveyed in yeast using microarray^27^, the determinants of NAC interaction with nascent chains (NCs) and its function beyond ER targeting and N-terminal processing are unknown. In particular, how NAC globally contributes to cotranslational proteostasis remains poorly understood (Fig. 1a).

**Figure 1.**
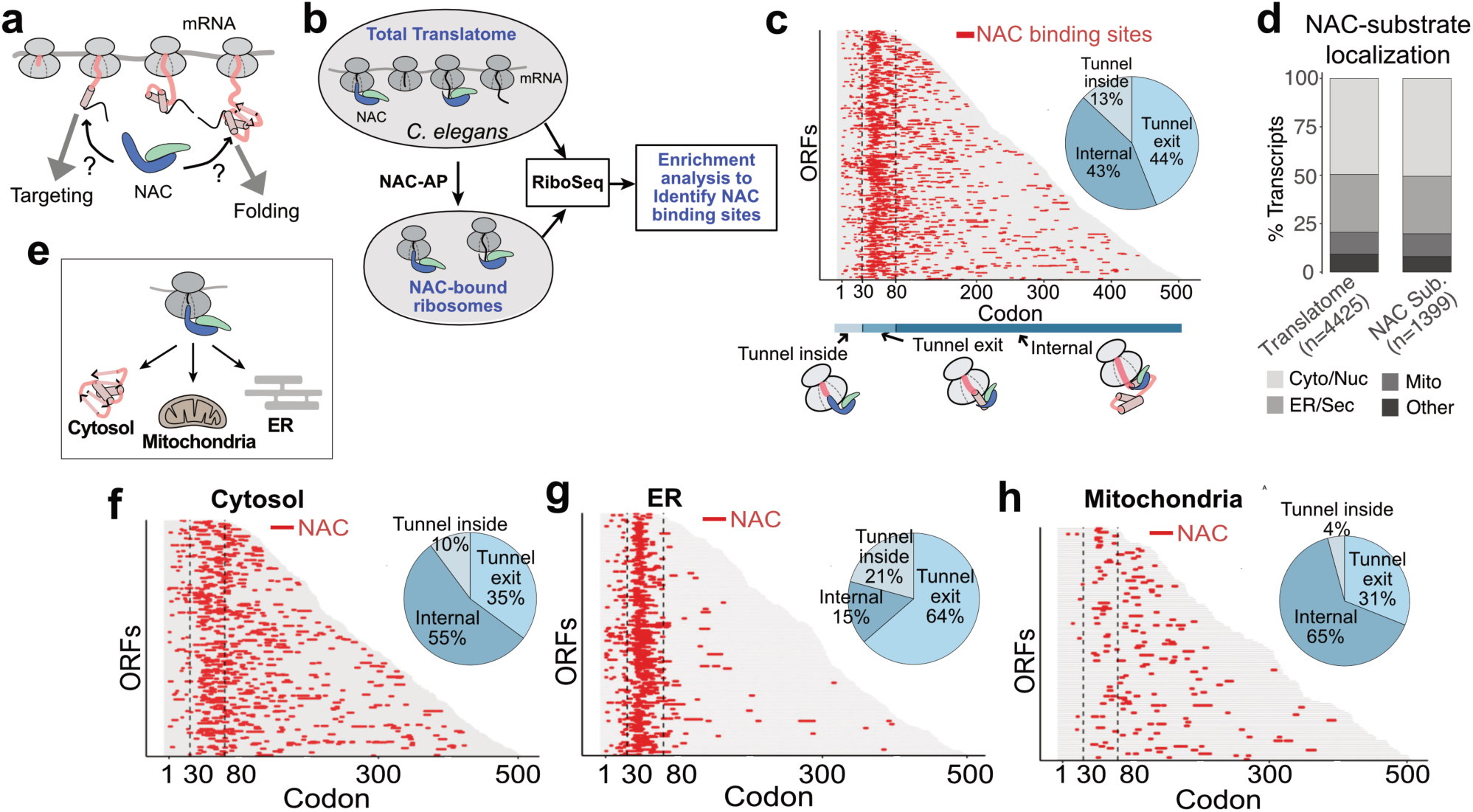
NAC interacts with broad range of nascent proteins throughout translation. **a,** Outline of questions on the role of NAC in protein biogenesis. **b,** Schematic of NAC-selective ribosome profiling and analysis^2^ (see Methods). AP: Affinity Purification. **c,** Positions of NAC interaction with nascent proteome (*red*) in three regions: Tunnel-inside (codon <30), Tunnel-exit (30 < codon < 80), and Internal (codon>80). **d,** Localization of proteins in translatome (transcripts analyzed) and identified NAC substrates. **e**, Schematic on how NAC may function distinctly on substrates destined to different compartments. **f-h,** Average enrichment of NAC and positions of NAC interaction with proteins in cytosolic/nuclear (**f**), ER/secretory (**g**), or mitochondria (**h**).

Here we integrate selective ribosome profiling^14^ and biochemical methods to define its critical proteostasis functions across all stages of translation. We show NAC cotranslationally protects hydrophobic helices in cytonuclear, ER, and mitochondrial nascent chains, ensuring their proper folding and transport. Remarkably, we discover an unexpected intra-tunnel sensing mode, where NAC engages ribosomes with extremely short nascent polypeptides inside the exit tunnel in a sequence-specific manner. These early NAC interactions tune ribosome flux during early synthesis in a phenomenon previously described as the elongation ramp^31–33^. We conclude NAC plays a pivotal role as a chaperone, targeting factor and regulator of translation kinetics during protein biogenesis.

### NAC interacts with a broad range of substrates at different nascent chain lengths

To globally map cotranslational interactions between nascent proteins and NAC, we performed NAC-selective ribosome profiling^14^. NAC-bound ribosomes were isolated from *C. elegans* through Twin-Strep tag (Extended Data Fig. 1a-e), and ribosome footprints were sequenced (Fig. 1b). The affinity tag did not affect NAC interaction with ribosomes (Extended Data Fig. 1c-d) and replicates of sequenced data were well correlated at both transcript and codon levels (Extended Data Fig. 1f-i). Moreover, ribosome-protected fragment sizes from all samples showed very good periodicity (Extended Data Fig. 1k) confirming that sequenced fragments are from translating ribosomes. Transcripts with good coverage and replicate correlation (n=4425) were analyzed for positional enrichment in NAC-bound ribosome footprints compared to the total translatome^2^ (see Methods). This allowed the identification of 2578 unique NAC binding events in 1399 proteins (Extended Data Fig. 1j). NAC interacts cotranslationally with proteins destined to both cytosol and nucleus (Cyto/Nuc) as well as ER and mitochondria (Fig. 1d), indicating a widespread and versatile role during biogenesis of proteins from distinct cellular compartments. Mapping the NAC binding sites along the open reading frame sequences (ORFs) revealed three types of NAC interactions with nascent chains (NCs). One class of binding events occurred when ∼50-60 residues of the NC were synthesized (Fig. 1c), which corresponds to the length at which NCs just emerge out of the ribosome exit tunnel. This is consistent with the previously suggested function of NAC as the factor that engage NCs as they first exit the ribosome^23,28–30^. Importantly, many NAC binding events occur at later, “internal” positions throughout the NC, indicating that NAC also binds NCs later during elongation. Surprisingly, we also observed a class of NAC binding events before the NC is exposed outside of the exit tunnel (Fig. 1c). To quantify the modes of NAC-NC binding, we divided the binding events into three regions: “*Tunnel-inside”* for NCs less than 30 aa in length, which have not yet emerged from the exit tunnel; “*Tunnel-exit”*, for NCs between 30∼80 aa which just emerged from the tunnel; and “*Internal”* for binding events occurring on NCs over 80 aa. NAC binding events in *Tunnel-exit* and *Internal* regions each represented roughly 44% of binding sites, while 13% of binding events occurred in *Tunnel-inside* (Fig. 1c). This pronounced interaction of NAC with ribosomes carrying very short NCs is in stark contrast with the binding mode observed for other cotranslationally acting chaperones like Hsp70 and TRiC, which associate with nascent chains after they emerge from the ribosome exit tunnel^2,34–36^.

We next examined how NAC engages substrates destined to different cellular compartments (Fig. 1e), including cytonuclear (Cyto/Nuc), mitochondrial (Mito) and ER/secretory (ER/Sec) proteins. We observed compartment-specific differences in the timing of NAC recruitment to NCs. Cytonuclear proteins associated with NAC throughout translation with a small bias towards the *Internal* binding mode (Fig. 1f). NAC interactions with mitochondrial proteins showed a strong bias towards *Internal* region binding and virtually no interactions in the *Tunnel-inside* region (Fig. 1h). In contrast, for ER and secretory (ER/Sec) proteins, NAC interactions were highly biased towards early binding events in *Tunnel-exit* and *Tunnel-inside* region (Fig. 1g). This mode of ER/Sec association is consistent with NAC acting prior to the delivery of ribosome-nascent chain complexes (RNCs) to the ER via SRP^23^. The differences in the timing of NAC recruitment to RNCs translating polypeptides destined to different cellular compartments highlights our ability to capture *bona fide* interactions and suggest distinct functions of NAC based on the ultimate fate of the nascent protein being synthesized. In particular, the predominance of *Tunnel-exit* NAC interactions with ER/Sec proteins supports an involvement of NAC in cotranslational targeting, while the large fraction of *Internal* NAC interactions with Cyto/Nuc and Mito proteins is consistent with a global chaperone function of NAC during cotranslational folding and maturation of nascent proteins. Of note, some NAC substrates exhibited multiple NAC recruitment events, suggesting NAC can re-engage *Internal* sites in the NC during synthesis (Extended Data Fig. 2a).

### NAC recognizes hydrophobic and amphipathic helical motifs in cytonuclear proteins

We next examined the role of NAC in cytonuclear protein biogenesis. NAC binding sites were enriched in proteins belonging to specific domain families, suggesting a chaperone role for these domain folds. Two different domain classification systems (PFAM^1^ and CATH^3,37^) revealed that specific domains including histones, RNA binding motifs, and nucleoside triphosphate hydrolases (NTPases) were enriched for NAC binding (Fig. 2a). To investigate the link between NAC binding and folding of specific domains, we mapped NAC binding sites around protein domain boundaries (Fig. 2b). We observed that NAC preferentially binds when ∼40 residues of a domain is exposed outside of the exit tunnel (Fig. 2c), suggesting a chaperone role stabilizing cotranslational domain folding intermediates.

**Figure 2.**
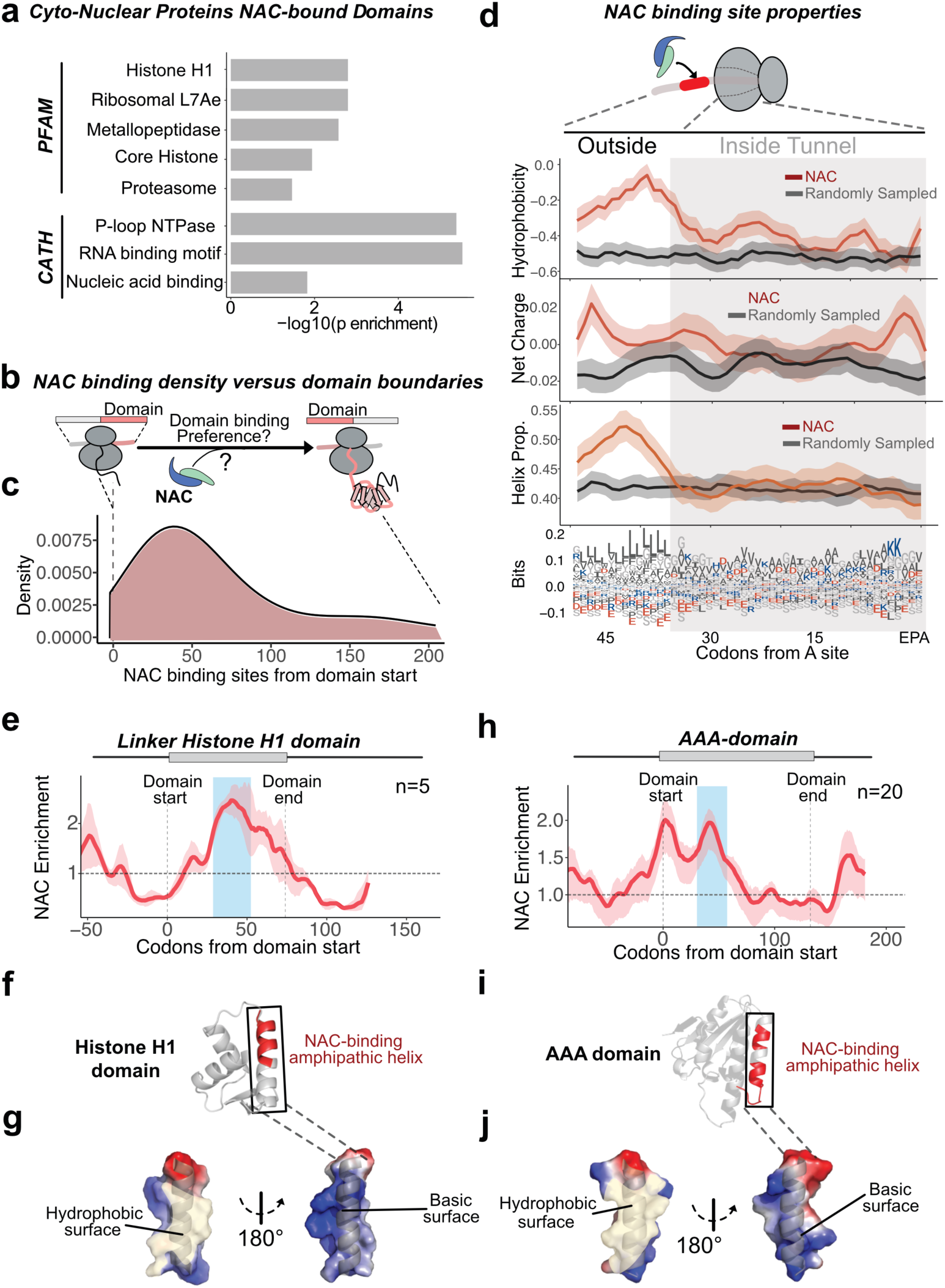
NAC recognizes hydrophobic helices to chaperone specific emerging domains. **a,** Enrichment of protein domains with NAC binding sites within the domain boundary. PFAM^1^ and CATH/Gene 3D^3,4^ classification systems were used. p value for enrichment was calculated using Fisher’s exact test comparing occurrence of domains in the NAC binding domains to the occurrence in the proteome. **b,** Schematic on whether NAC interacts within the boundaries of protein domains. **c,** Distribution of NAC binding positions from the start of the domains. **d,** Average of physiochemical properties of nascent chains aligned to NAC binding sites in cytonuclear proteins within both Tunnel-exit and Internal regions. The shaded region represents a region buried inside the ribosome exit tunnel that spans 35 amino acids of nascent chains. **e-j,** Examples of average NAC enrichment in Linker Histone H1 domain (**e-g**) or AAA domain (**h-j**). Full-length predicted structures^7,8^ of each domain (**f** and **i**) and surface representation of NAC-interacting helix (**g** and **j**) are shown with NAC binding sites highlighted in red. The enrichment plot and NAC binding sites were offset by 35 codons to depict the region that is exposed outside of the exit tunnel when NAC binds.

We next asked if NAC recognizes a specific motif in its cytonuclear substrates during synthesis (Fig. 2d). When we aligned all NAC binding sites on cytosolic substrates and analyzed physiochemical features of the nascent chains emerging from the ribosome upon NAC binding (Fig. 2d), we found the region just outside the exit tunnel (∼40 codons from A-site) was highly hydrophobic and preceded by a positively charged patch. Additionally, this region had high helical propensity, showing that NAC generally recognizes hydrophobic segments with helical propensity emerging out of the ribosome exit tunnel (Fig. 2d). Importantly, these properties were observed for both *Tunnel-exit* and *Internal* NAC binding sites as they emerged from the ribosome, indicating these sequence features represent a chaperone-recognition motif for NAC regardless of its position along the transcript (Extended Data Fig. 2b and c). Furthermore, NAC could recognize these motifs multiple times during synthesis of a given transcript (Extended Data Fig. 2a), suggesting it re-engages the nascent chain once a suitable binding site is exposed. Of note, the NAC-recognized cytonuclear helical motifs exhibited higher amphipathicity compared to the more hydrophobic signal sequences or transmembrane domain motifs recognized by SRP (Extended Data Fig. 2d).

We also compared the sequence length of the NAC binding motifs identified in our analysis with those recognized cotranslationally by another chaperone, the Hsp70 SSB, in yeast, which is known to bind to hydrophobic regions with beta-sheet propensity^2^. Despite their distinct recognition, both NAC and Hsp70 bind motifs of similar width (Extended Data Fig. 2e), suggesting that NAC is a *bona* fide cotranslational chaperone non-redundant to Hsp70.

To further investigate the link between NAC binding events and folding of specific domains, we mapped the NAC binding sites on the sequence of two domain substrates, the linker Histone H1 and the AAA-ATPase domains. Metagene analysis of all histone H1 domains (n=5) showed a shared pattern of prominent NAC enrichment when ∼40 residues of the domain were exposed (Fig. 2e), which is when the third helix is just emerging from the ribosome exit tunnel (Fig. 2f). Similarly, metagene analysis of the AAA-ATPase family domains (n=20) showed prominent enrichment of NAC (Fig. 2h) during synthesis of the second helix (Fig. 2i). Interestingly, both NAC-binding helices of H1 and AAA domains showed amphipathic character (Fig. 2g and j), with the hydrophobic side facing the structural core, while the basic surface faces the solvent. Our data suggests that NAC preferentially interacts with hydrophobic or amphipathic helices emerging from the ribosome and protects their exposed hydrophobic side until further translation of the domain allows them to get buried into the structural core of the domain.

Together, these results suggest that NAC is a chaperone that binds hydrophobic helices of a wide range of protein domains, especially nucleic acid interacting domains, to assist in their cotranslational domain-wise folding. Importantly, this sets NAC apart from cotranslational Hsp70 chaperones, which recognize hydrophobic beta-sheets^2,36^, suggesting a nonredundant role of NAC in cotranslational folding. The similar binding width of NAC and Hsp70 (Extended Data Fig. 2e) suggests a transient and dynamic interplay between these chaperones on the nascent chains on the ribosome.

### NAC chaperones a subset of mitochondrial precursors

Little is known about NAC function in mitochondrial import, although a potential role in import of some mitochondrial proteins was previously suggested^38–44^ (Fig. 3a). Our analysis showed many nuclear encoded mitochondrial proteins interact cotranslationally with NAC (Fig. 1h), including mitochondrial precursors destined to the mitochondrial matrix, outer membrane (OM), and inner membrane (IM) (Extended Data Fig. 3a). Further analysis showed minimal overlap between NAC binding and the location of a N-terminal mitochondrial targeting sequence (MTS), suggesting NAC does not act as an MTS binding and targeting factor (Fig. 3b). On the other hand, we observed significant overlap between NAC binding sites and transmembrane helices (TMs) of mitochondrial membrane proteins (Fig. 3c and Extended Data Fig. 3b-c), particularly for early emerging TMs. This suggests that NAC protects these hydrophobic domains to maintain the nascent protein competent for import into the mitochondrial membrane. To better understand the possible function of NAC in mitochondrial protein biogenesis, we mapped NAC binding sites to their mature protein structures. Consistent with the analyses of NAC binding for cytonuclear proteins, NAC binding sites (highlighted in red) in mitochondrial precursors also mapped to helical regions and TMs motifs as seen in membrane proteins such as SLC-25A10, MTCH-1, and TSPO-1 (Fig. 3d and Extended Data Fig. 3e-f) as well as for membrane proteins in Complex II of the respiratory chain (Fig. 3e and Extended Data Fig. 3d). Interestingly, NAC binding regions also mapped to motifs that will become hydrophobic cofactor binding pockets, as highlighted for the FAD, UQ1 and heme binding sites in Complex II (Fig. 3e), suggesting that NAC protects these hydrophobic pockets from aggregation to maintain the NC in an import-competent state.

**Figure 3.**
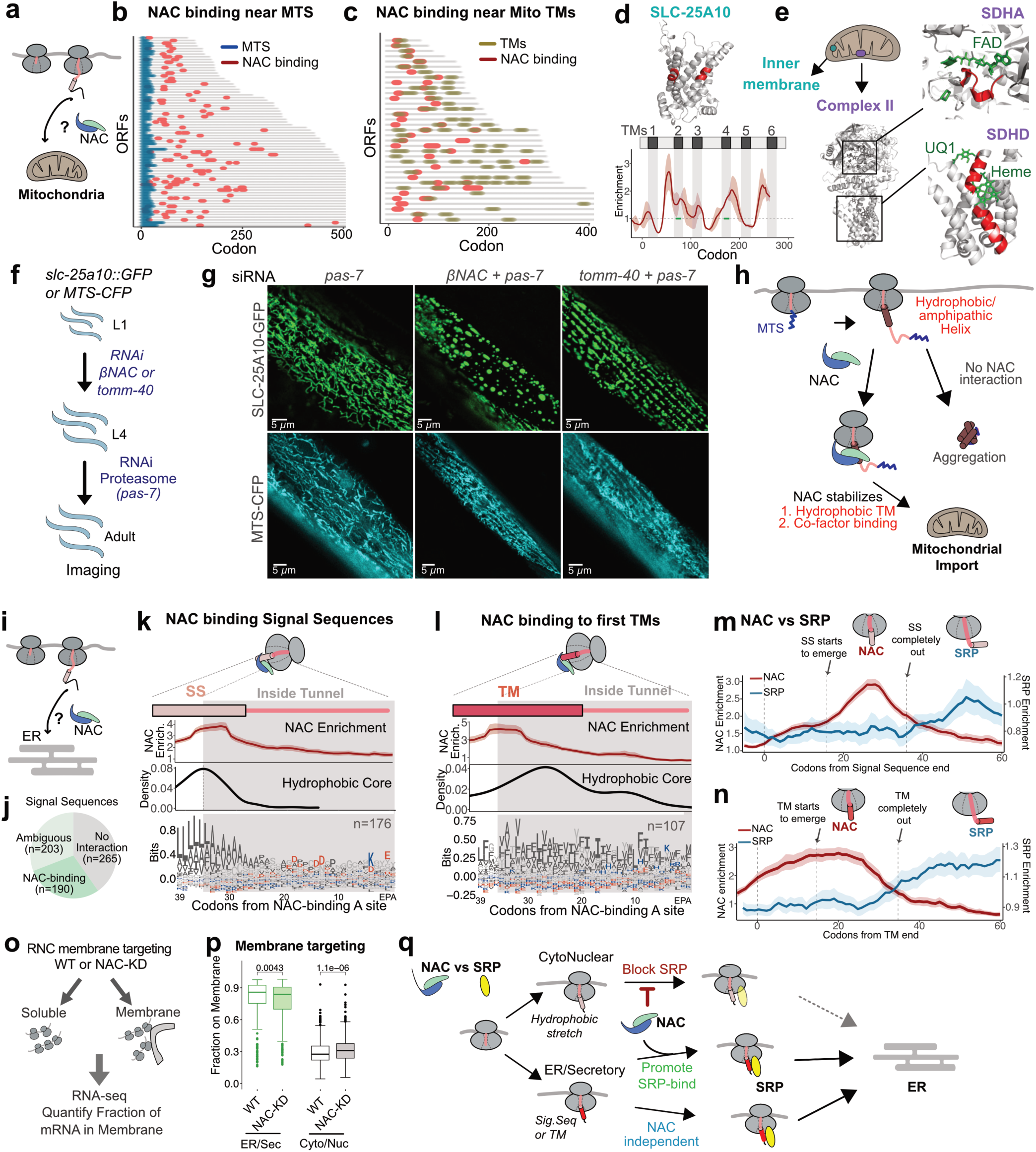
NAC plays an essential role during the biogenesis of mitochondrial and ER proteins. **a,** Outline of investigation on the role of NAC in biogenesis of mitochondrial proteins. **b,** ORFs with MTS (blue) and NAC binding sites (red). Common predictions from MitoFates^5^ and TargetP^6^ were taken as MTS. **c,** ORFs with transmembrane helices (TM, light brown) and NAC binding sites (red). Only TMs annotated in Uniprot were used. **d-e,** Examples of mitochondrial inner membrane proteins of SLC-25A10 (**d**) and Complex II (**e**) with NAC binding sites (red) within TMs. Structures predicted from Alphafold^7,8^ were overlayed to available structures 8GS8^9^ to show cofactor (green) binding. **f,** Schematic to investigate how NAC or tomm-40 knockdown affects mitochondrial protein import in C. elegans. Knockdown of NAC or tomm-40 was introduced at L1 stage and pas-7 knockdown at L4 stage before imaging. **g,** Microscopy of SLC-25A10-GFP (top) and MTS-CFP (bottom) with knockdown of pas-7 (left), NAC/pas-7 (middle), or tomm-40/pas-7 (right) shown in single muscle cells of C.elegans. **h,** Summary of the role of NAC in mitochondrial protein biogenesis. While NAC does not interact directly with MTS, it protects hydrophobic TMs during transit to mitochondria. Failure to interact with NAC leads to aggregation. **i,** Outline of investigation on the role of NAC during the biogenesis of ER-destined proteins. **j,** Fraction of ER-localized proteins with N-terminal signal sequence that interact with NAC during synthesis. Ambiguous group represents proteins that show some degree of NAC enrichment but not enough to pass the threshold (see Methods). **k-l,** Sequence features of SS (**k**) or TMs (**l**) aligned to NAC binding sites. The shaded region represents a region buried inside the ribosome exit tunnel that spans 35 amino acids of nascent chains. NAC enrichment plot was offset by 35 amino acids to show where it interacts with exposed nascent chains. **m-n,** Enrichment of NAC and SRP aligned to the end of signal sequences (**m**) or TMs (**n**). SRP data was taken from^15^. **o,** Schematic of membrane fractionation and RNA-seq to quantify the extent of cotranslational targeting in WT or with NAC-KD. **p,** The fraction of ER/Sec (green) (n=366) or Cyto/Nuc (grey) (n=624) mRNAs localized on the membrane with or without NAC-KD for NAC substrates. Statistical analysis was performed using two-sided Wilcoxon rank-sum tests. **q,** Summary of NAC function in targeting ER-destined proteins. NAC prevents SRP engagement with hydrophobic cytosolic nascent chains while actively relaying a subset of SS/TMs to SRP.

We next assessed the role of the NAC-TM interactions in mitochondrial protein import *in vivo*. We monitored the mitochondrial localization of NAC substrate SLC-25A10 (Fig. 3d) bearing a C-terminal GFP in animals where NAC was depleted by knockdown of NACβ (Fig. 3f). As a control, we also examined an MTS-bearing soluble mitochondrial-targeted CFP (MTS-CFP), which should be imported independently of NAC (Fig. 3f). To stabilize the potentially non-imported SLC-25A10 membrane protein from cytosolic degradation, we also inhibited the proteasome system by knockdown of the proteasome subunit PAS-7 (Fig. 3f, 3g and Extended Data Fig. 3g). While *pas-7* knockdown alone did not affect the mitochondrial localization of SLC-25A10-GFP (Fig. 3g, *left*), co-knockdown of NACβ induced formation of punctate, presumably aggregated, structures of SLC-25A10-GFP (Fig. 3g, *middle*). The mitochondrial localization of MTS-CFP was unaffected by NAC knockdown (Fig 3g, bottom panel). A similar albeit less pronounced punctate pattern of SLC-25A10-GFP was also observed by knockdown of *tomm-40*, a mitochondrial protein import receptor^45^ (Fig. 3g, *right*). Visualization of mitochondria using a fluorescent dye (Mitotracker) in NAC and tomm-40 knockdown backgrounds revealed partial disruption of the fine mitochondrial network; however, overall mitochondrial integrity was largely preserved (Extended Data Fig. 3i). These observations indicate that NAC contributes to maintaining general mitochondrial integrity, but the aggregation phenotypes observed with SLC-25A10-GFP are specific to NAC-dependent mitochondrial substrates. Together, these data suggest that NAC serves as a cotranslational chaperone that protects hydrophobic motifs in mitochondrial precursors, including TMs and cofactor binding pockets, to prevent their aggregation and allow proper import into mitochondria (Fig. 3h).

### Defining the multifaceted role of NAC in ER targeting

Previous studies showed that NAC cooperates with SRP in cotranslational targeting of ER-destined proteins^23–25^. However, it remains unclear whether NAC interacts universally with all ER-destined proteins and how precisely the timing of these interactions is regulated. To answer these important questions, we examined NAC interactions with signal sequences (SS) and TMs of ER-destined proteins (Fig. 3i). As described above, NAC primarily interacts with ER-destined RNCs at the *Tunnel-exit* region (Fig. 1g) consistent with its role during pre-targeting of soluble RNCs. Notably, approximately 30% of ER-targeting SS/TMs associated with NAC (Fig. 4j and Extended Data Fig. 4a), suggesting that only a specific subset of ER-destined proteins are dependent on NAC. To identify features dictating the specificity of NAC interactions, we compared the sequence properties of ER-destined RNCs that either do or do not interact with NAC. NAC preferentially interacts with more hydrophobic signal sequences, composed of less bulky amino acids and containing a positively charged N-terminus (Extended Data Fig. 4b-d). For those TMs that did bind NAC, there was no preference for hydrophobicity. However, as observed for SS, NAC preferred TMs with less bulky residues and a slight preference for a positively charged N-termini (Extended Data Fig. 4b-d).

**Figure 4.**
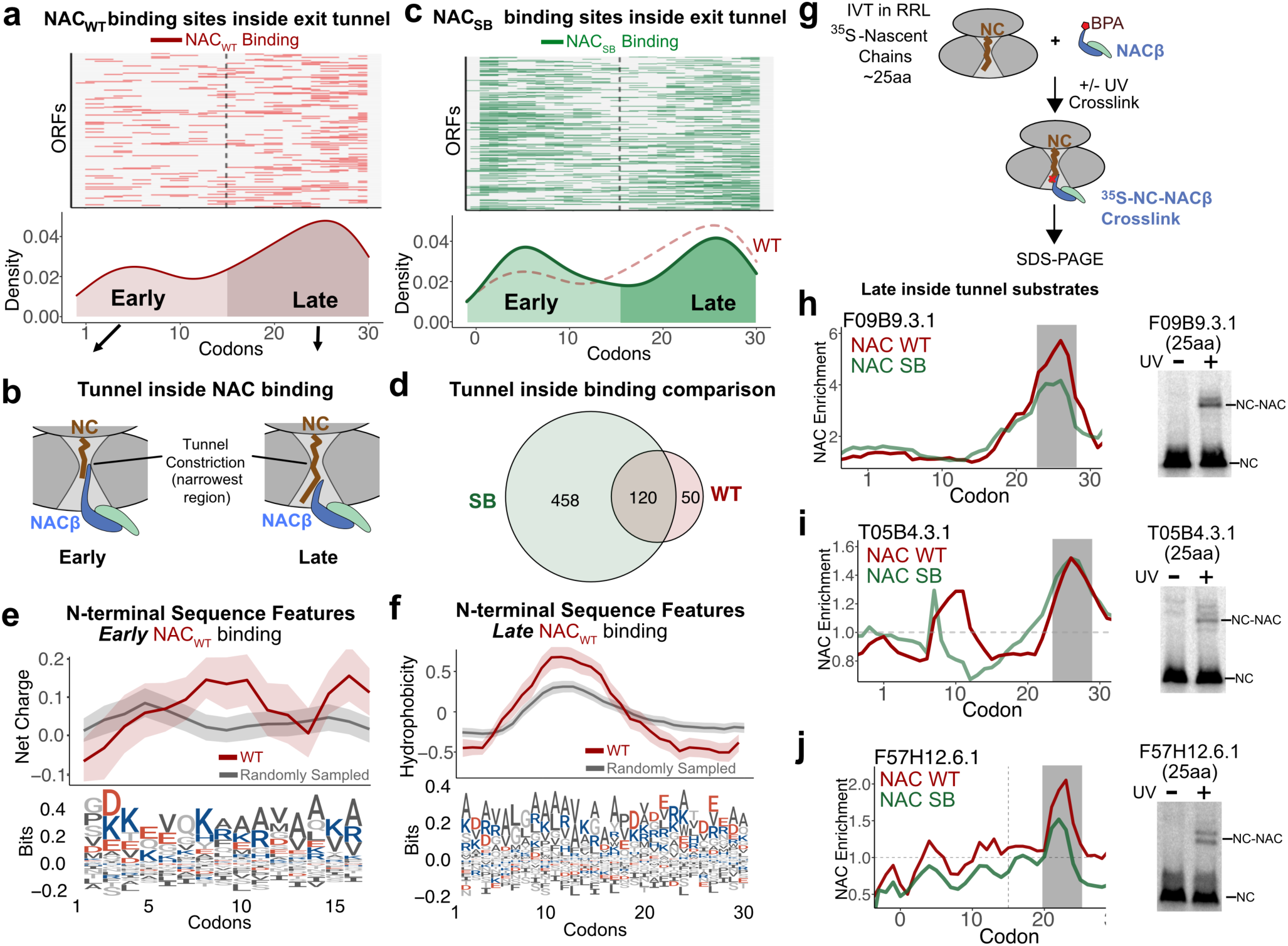
NAC interacts with NCs inside the ribosome exit tunnel by recognizing specific sequence features. a, Distribution of NAC_WT_ binding sites during the synthesis of the first 30 amino acids. **b,** Schematic of NAC interactions with NC inside ribosome exit tunnel. NAC binding to NC lengths 1-15aa is defined as ‘**Early** binding’ and NAC binding to NC lengths 16-30 aa is defined as ‘**Late** binding’. **c,** Distribution of NAC binding sites of ribosome super-binder NAC_SB_ during the synthesis of the first 30 amino acids. **d**, Tunnel inside interacting substrate comparison between WT and SB NAC. **e,** Sequence features of N-termini of NC that interact with NAC during the incorporation of first 15 amino acids (**Early**, *red*) or randomly sampled (*grey*). **f,** Sequence features of N-termini of NCs that interact with NAC during the synthesis of15 to 30 amino acids (**Late**, *red*) or randomly sampled (*grey*). **g**, Schematic of crosslinking experiment. Ribosome-nascent chain complex bearing ∼25 aa NC are synthesized with [^35^S]Met, incubated with Bpa-NAC, exposed to UV for crosslinking, and resolved on SDS-PAGE^10^. **h-j**, Products with or without UV resolved on SDS-PAGE and NAC enrichment plot for corresponding substrates, F09B9.3.1 (**h**), T05B4.3.1 (**i**), and F57H12.6.1 (**j**). The lower band represents the non-crosslinked NC, and the upper band represents the crosslinked product between the NC and NAC (NC-NAC).

To gain insight into the temporal regulation of the NAC interaction vis-a-vis SRP recruitment during the translation of SS/TMs, we analyzed the sequence features surrounding the NAC binding sites. NAC interacts with secretory RNCs when SS/TMs are only partially exposed outside of the exit tunnel (Fig. 3k-l). NAC binds SS when the most hydrophobic part starts to emerge from the exit tunnel and with TMs just before the most hydrophobic region is exposed outside the exit tunnel (Fig. 3k-l). Exploiting the conservation of SRP across species^46,47^ we compared the timing of NAC and SRP engagement, by analyzing SRP selective ribosome profiling data from yeast^15^. Comparing NAC and SRP enrichment upon emergence of SS/TMs outside the exit tunnel, we observe that NAC binds much earlier than SRP (Fig. 3m-n, *red*). On the other hand, only SRP engages the SS/TMs once they completely emerge outside of the exit tunnel (Fig. 3m-n, *blue*). While this comparison is done using data from two different organisms, these experiments suggest that NAC and SRP engage SS/TMs with distinct and sequential timing as they emerge from the ribosome exit tunnel. These findings are consistent with the proposed NAC to SRP handover model inferred from cryoEM structures of NAC-RNC complexes^23^.

We next tested whether NAC binding to RNCs bearing SS/TMs are important for efficient targeting. We performed membrane fractionation followed by RNA-Seq in wildtype (WT) and NAC knockdown (NAC-KD) worms and quantified membrane-bound mRNA, which reports on the degree of cotranslational membrane targeting (Fig. 3o). Soluble and membrane fractions were isolated reproducibly with minimal cross-contamination (Extended Data Fig. 4e-g). Importantly, NAC depletion (NAC-KD) did not affect membrane targeting of ER/Sec proteins that do not interact cotranslationally with NAC (Extended Data Fig. 4h). This indicates that not all SS/TM-containing proteins require NAC for proper ER targeting. In contrast, those ER/Sec proteins that do interact cotranslationally with NAC showed a mild but significant reduction in membrane targeting (Fig. 3p, *green*). This trend was maintained when we quantified the targeting defect for SS/TM-containing proteins separately (Extended Data Fig. 4i). We conclude that NAC specifically engages a subset of SS/TM-containing proteins to selectively enhance their membrane targeting. Of note, NAC-KD led to mistargeting of some cytonuclear proteins to the membrane (Fig. 3p, *grey* and Extended Data Fig. 4h). These analyses highlight the nuanced role of NAC in ER targeting; NAC prevents mistargeting of Cyto/Nuc proteins to the membrane while selectively enhancing membrane targeting of ER-destined proteins by cotranslational interactions with SS/TMs as they emerge from the ribosome (Fig. 3q).

### NAC selectively interacts with nascent chains inside the ribosomal tunnel

The narrow ribosome exit tunnel protects ∼30-40 residues of the nascent chain^18,48^. All cotranslational acting chaperones analyzed to date^2,15,34–36^, associate with the nascent chains at lengths over ∼30-40 residues, i.e. once the nascent chains exit the ribosome. However, we observe N-terminal NAC enrichment on very short nascent chains (<30aa) that are inside the ribosome exit tunnel (Fig. 1c, Tunnel inside). Intriguingly, mapping all *inside-tunnel* NAC interactions revealed a bimodal NAC binding pattern, with NAC binding early (average around 5 aa) or late (average around 25 aa) within the tunnel (Fig. 4a, left). We rationalized this observation by considering that the ribosome exit tunnel is not uniform^48,49^. A constriction defines two regions based on where the emerging N-terminus is located: the upper region proximal to the PTC (NC 1-15 aa, herein ‘**Early**’) (Fig. 4b, left) and the lower region past constriction to the tunnel exit (NC 15-30, herein ‘**Late**’) (Fig. 4b, right). Interestingly, previous structural and biochemical data indicated that the flexible N-terminus of NACβ can insert deep into the tunnel, reaching near the constriction site^10^. This raises the possibility that NAC contacts a subset of nascent chains while they are still inside the tunnel via the N-terminus of NACβ (Fig. 4b).

To further assess NAC interactions with nascent chains inside the exit tunnel, we exploited previous findings indicating that the insertion of the NACα N-terminus into the tunnel is enhanced in a NAC mutant (ΔN1-53 NAC) lacking the flexible NACα N-terminus which acts as a negative regulator of NAC-ribosome binding^10^. Accordingly, ΔN1-53 NAC displays an increased affinity for ribosomes. We predicted that this higher affinity ΔN1-53 NAC mutant (herein Super-Binder NAC or SB-NAC) should stabilize the NAC-nascent chain interactions. NAC-selective ribosome profiling experiment in animals/C.elegans expressing SB-NAC replicated our findings for WT NAC (Extended Data Fig. 5). The SB-NAC and WT-NAC interaction data were well correlated at transcript and codon levels (Extended Data Fig. 5a-d) and identified similar substrates overall (Extended Data Fig. 5e-j). Consistent with our prediction, we identified significantly more NAC-nascent chain interactions for SB-NAC, particularly for binding events inside the exit tunnel, including both **Early** and **Late** interactions (Fig. 4c, 4d, and Extended Data Fig. 5k-m). **Early** tunnel interactions were particularly enriched in SB-NAC over WT-NAC (Fig. 4c, bottom panel).

Since only a subset of NCs exhibit intra-tunnel NAC-interactions, we next examined if there was a sequence specificity driving in tunnel NAC association. Indeed, both the **Early** and the **Late** intra-tunnel NAC interaction displayed distinct NC sequence features. The N-termini of NAC substrates with **Early** intra-tunnel interactions were generally highly charged with a bias towards negative charge at the very N-terminus (Fig. 4e). On the other hand, the N-termini of NAC substrates with **Late** intra-tunnel interactions were hydrophobic enriched in Alanine and aliphatic residues (Fig. 4f). Importantly, the NAC-nascent chain recognition specificity was the same for both WT-NAC and SB-NAC. **Early** nascent chain interactions of SB-NAC also contained charged N-termini (Extended Data Fig. 5n), and **Late** interactions also contained hydrophobic N-termini (Extended Data Fig. 5o). The consistent results between these two NAC variants strongly suggest that we are capturing bona fide *in vivo* NAC-nascent chain interaction inside the exit tunnel. Furthermore, the observed sequence specificity strongly suggests NAC is directly interacting with a subset of the nascent chains very early during translation when NCs are still inside the tunnel.

To verify that NAC interacts with NCs within the tunnel, we used in vitro translation (IVT) to generate short RNCs bearing ^35^S-labeled 25 aa-long nascent chains corresponding to three NAC substrates identified *in vivo* and having a **Late** *inside-tunnel* interaction (F09B9.3.1, T05B4.3.1, and F57H12.6.1). These RNCs were incubated with recombinant NAC carrying a photo-crosslinking amino acid (Bpa) at the first position of NACβ^10,23^ (Fig. 4g). For each of these RNCs, we observed crosslinking between NAC and the ribosome-bound ^35^S-labeled NAC substrate (Fig. 4h-j), demonstrating a direct interaction between NACβ and the nascent chain inside the exit tunnel (‘**Late**’ region). To further validate that these interactions depend on the insertion of the N-terminus of NACβ into the exit tunnel, we generated a NAC variant where a ∼100 aa globular SUMO domain is fused at the N-terminus of NACβ. The well folded SUMO domain should prevent the N-terminus of NACβ from inserting into the exit tunnel. Indeed, none of the tested substrates crosslinked to SUMO-NAC confirming that this interaction depends on NACβ insertion into the exit tunnel (Extended Data Fig. 6). Interestingly, AlphaFold (AF) modeling offers a rationalization of how NACβ in the exit tunnel could associate with the nascent chain in the wider vestibule region of the exit tunnel (Extended Data Fig. 5q-r). Thus, AF predicts the N-terminus of NACβ can form an amphipathic helix that interacts with the hydrophobic NC **Late** intra-tunnel segment (Extended Data Fig. 5q-r). Together, these results show a novel mode of NAC interaction with nascent chains inside the ribosome exit tunnel.

### Intra-tunnel NAC interactions with nascent chains modulate translation elongation

Interactions between the nascent chain and the exit tunnel wall may lead to a slowdown of elongation^18,50,51^. We thus considered whether intra-tunnel interactions between NAC and NCs may affect their translation speed (Fig. 5a). To test this hypothesis, we carried out ribosome profiling^52^ analysis comparing WT and NAC-depleted worms (NAC-KD) (Fig. 5a and Extended Data Fig. 7a-b). Metagene analysis of WT animals revealed previously described early elongation slowdown events before the NC first emerges from the ribosomal tunnel, which reflect the slow elongation kinetics early in translation^31,33,53,54^ (Fig. 5b-c, N2(WT)). These include an early accumulation of ribosome density around codon five, proposed to represent a slowdown of the NC near the constriction site^33^. We also observed the increased ribosome density between residues 5-30 previously termed the ‘ramp’, proposed to tune translation efficiency^31^ (Fig. 5b-c, N2(WT)). Strikingly, these early slowdown events were nearly completely abrogated upon NAC knockdown (Fig. 5b-c, NAC-KD). The peak around codon five became much lower, while the ‘ramp’ was no longer detected (Fig. 5b-c, NAC-KD). These results indicate that these early elongation slowdowns, including the previously described “ramp”, are mediated by NAC.

**Figure 5.**
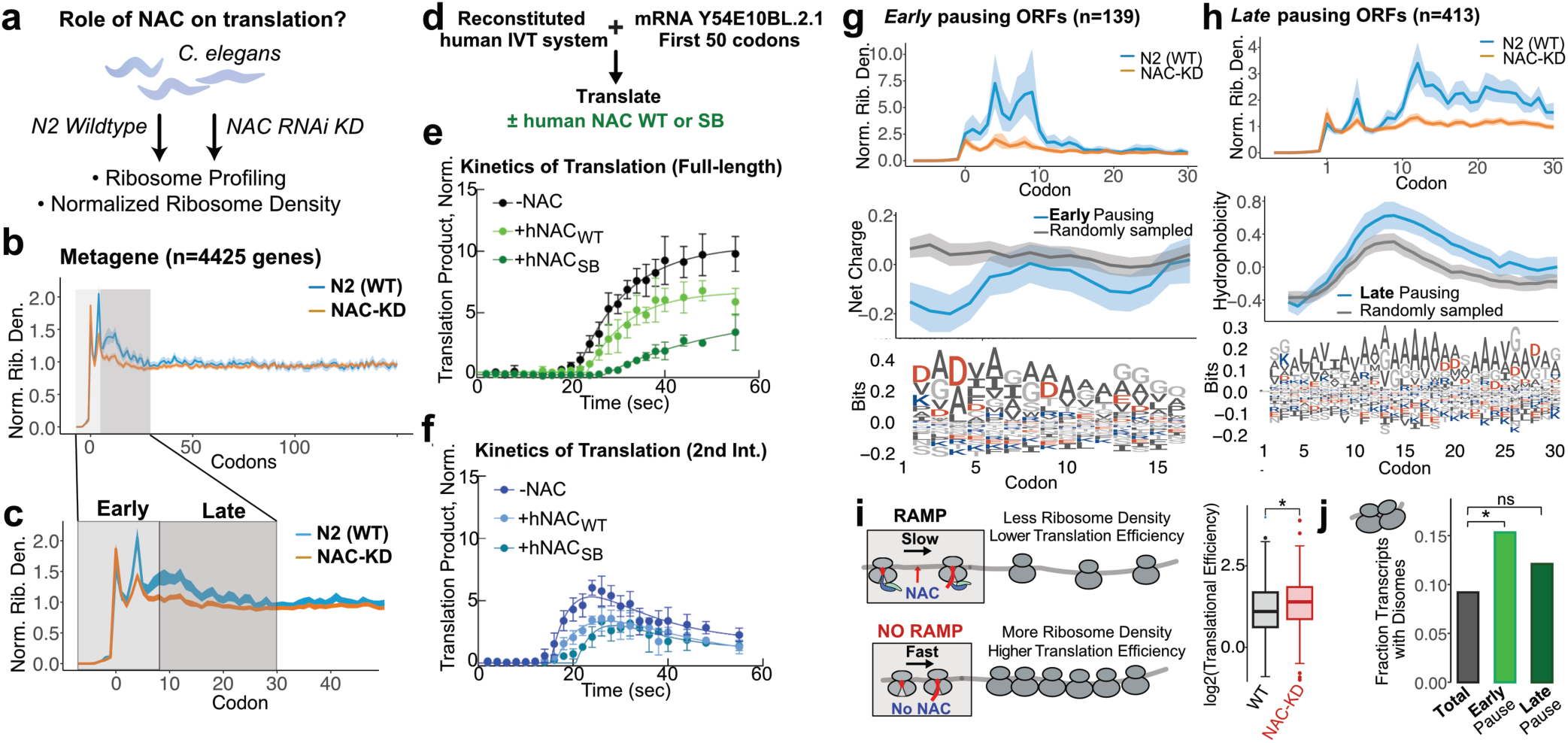
Tunnel-inside NAC interactions with RNCs induce translation slowdowns. **a,** Schematic of ribosome profiling from WT or NAC-KD worms to investigate role of NAC in modulating translation speed. **b,** Metagene (n=4425) profile of average ribosome density around start codon from WT (*blue*) or NAC-KD (*orange*) worms showing two distinct peaks near **Early** and **Late** regions. **c,** Same as in **b** but zoomed in to first 50 codons. **d,** Schematic of *in vitro* translation experiment. The first 50 residues of Y54E10BL.2.1 are translated in a synchronized reconstituted eukaryotic IVT with initiator BODIPY-Met-tRNA in the absence or presence of NAC, quenched at different time points, and translation products are resolved on Tris-Tricine SDS gel visualized through fluorescence imaging. **e-f,** Appearance of full-length translation product (**e**) or a translation intermediate (**f**), normalized to loading control in the absence (**e**:*black,* **f**:*blue*), in the presence of wildtype human NAC (**e**:*light green,* **f**:*skyblue*), or mutant human NAC_ΔN1-67_ (equivalent mutation to NAC_ΔN1-53_ in C. elegans, SB) (**e**:*dark green,* **f**:*teal*). **g-h,** Characteristics of transcripts with **Early** (**g**) or **Late** (**h**) pausing; Average ribosome density around start codon (top), net charge of N-termini (middle), and sequence logo of N-termini (bottom). **i**, A model of how early ramp may control the ribosome density on mRNA (left) and translation efficiency of transcripts with NAC-dependent slowdown in pre-constriction region (n=139, p=8.9x10^-4^) for WT and NAC-KD condition (right). Statistical analysis was performed using two-sided Wilcoxon rank-sum tests. **j,** Disome peaks that occur ∼10 residues from pause sites (pause score >6) were searched (see Methods) and the fraction of transcripts that have this peak were quantified and compared (Total: Disome (n=406)/All (n=4417), Early: Disome (n=21)/All (n=137), Late: Disome (n=52)/All (n=430)). Statistical analysis was performed using Fisher’s exact test. p=0.024 (Early) and p=0.057 (Late).

To directly test whether NAC regulates the rate of early elongation as the nascent chain first moves through the exit tunnel, we established a fully reconstituted eukaryotic *in vitro* translation (IVT) system using purified human components^55^ (Fig. 5d). As this IVT system lacks any chaperone, it allows us to directly investigate the role of NAC on individual steps of translation. To capture the effect of NAC in early translation, we translated only the first 50 codons of the endogenous NAC substrate col-48 (Extended Data Fig. 7c, Y54E10BL.2.1), which exhibits intra-tunnel NAC binding and exhibits a NAC dependent translation slowdown (Extended Data Fig. 7c). The reconstituted IVT system achieved an elongation rate of ∼2 aa/s based on the time when the full-length translation product is observed (Extended Data Fig. 7e), which is close to the reported rate of elongation *in vivo*^56^. This near-physiological rate of elongation allowed us to resolve fine details of translation, including two distinct translation intermediates during the synthesis of the N-terminal 50 aa-long peptide (Fig. 5d-f and Extended Data Fig. 7e-g, -NAC). Addition of purified wildtype human NAC slowed elongation and delayed the appearance of both intermediates, indicating that exogenous NAC can by itself lead to a slowdown in elongation (Fig. 5d-f, and Extended Data Fig. 7e-g, hNAC_WT_). Strikingly, the addition of the higher affinity human SB-NAC variant (ΔN1-67 NACα in human, equivalent to ΔN1-53 in *C. elegans*) caused an even stronger slowdown in elongation (Fig. 5d-f and Extended Data Fig. 7e-g, hNAC_SB_). Moreover, prolonged accumulation of the translation intermediates was observed in the presence of SB-NAC mutant (Fig. 5f and Extended Data Fig. 7f). These results show that purified NAC added to a reconstituted IVT system directly induces slowdown very early during the translation, consistent with our ribosome profiling data. The mechanistic underpinnings of this slowdown, i.e. whether it involves full or partial insertion of the intrinsically disordered N-terminus of NACβ into the tunnel, or a conformational change caused by binding to NAC to the ribosome will require further structural analyses. Our experiments indicate that the elongation slowdown is dependent on the strength of NAC interaction with the ribosome as evident with the increased slowdown upon SB-NAC addition, consistent with a previous IVT study using rabbit reticulocyte lysate (RRL)^10^. Addition of SUMO-NAC to the IVT, which cannot fully insert into the exit tunnel (Extended Data Fig. 6), still led to a slowdown in elongation, consistent with the effects of a tunnel-sensing mutant NAC variant in RRL that inhibited translation, albeit weaker than WT NAC^10^ (Extended Data Fig. 7h-k). These experiments show that full insertion of the N-terminus of NACβ is not required for translation slowdown. However, the disordered and highly charged NACβ N-terminus of SUMO-NAC could still partially loop into the tunnel, explaining the observed early slowdown of translation in the presence of NAC. Further structural analyses of this effect will be required to fully understand this mode of regulation.

To gain further mechanistic detail and explore the link between NAC-dependent early elongation slowdowns and NAC intra-tunnel interactions, we next identified any general sequence motifs in NCs whose translation elongation is influenced by NAC-KD. We analyzed transcripts that exhibit a NAC-dependent elongation slowdown, either early in the tunnel, i.e. around position five or late in the tunnel, i.e. at the ‘ramp’ (Extended Data Fig. 8a). Notably, the N-termini of transcripts subject to either mode of NAC-dependent slowdowns shared sequence features at the amino acid level rather than at the codon usage level, indicating these slowdowns are not driven by tRNA abundance (Extended Data Fig. 8b-c). Remarkably, the sequence properties for both “early” and “ramp” slowdowns resembled those of **Early** and **Late** NAC-binding motifs described above (compare Figs. 4e-f and 5g-h). Transcripts with NAC-induced pausing early in the tunnel contained a negatively charged N-terminus with aspartic acid enriched in the second and fourth position, resembling ‘**Early**’ intra-tunnel interactions of NAC (Fig. 5g and 4e). On the other hand, transcripts with NAC-dependent “ramp” pausing showed increased hydrophobicity (Fig. 5h and 4f), resembling ‘**Late’** intra-tunnel interactions of NAC. These similarity in features between the NAC-dependent slowdowns in the tunnel and the NAC binding motifs in the tunnel strongly suggest that translation slowdown events at the start of translation early and late in the tunnel are mediated by NAC-NC interactions.

The NAC-induced slowdown of elongation early in translation may serve several potential functions. For instance, the ‘ramp’ was speculated to controls ribosome flux through mRNAs to reduce ribosome density along the transcript^31^ (Fig. 5i, left). To check if this is the case for NAC-induced pausing, we quantified translation efficiency (TE), which reports on the relative density of ribosomes for a given transcript. Strikingly, NAC knockdown increased TE for transcripts that exhibit early NAC-induced pausing but not for those lacking a “ramp” (Fig. 5i, right). High ribosome density can lead to disome formation and increased risk of ribosome collisions. Notably, transcripts with a NAC-induced slowdown also contained a higher fraction of transcripts forming disomes (Fig. 5j). Of note, some of these pausing events were associated with ER membrane targeting (Extended Data Fig. 8d), showing important physiological role of NAC-induced pausing. Given the importance of elongation rate to additional protein biogenesis events, our data indicate that NAC directly modulates early elongation kinetics to optimize efficient translation to be attuned to protein targeting and folding.

## Discussion

Here we define NAC as a key cotranslationally acting factor that governs proteostasis, subcellular targeting, and translation elongation rate during protein biogenesis (Fig. 6). We show that NAC engages nascent chains even before they emerge from the ribosomal tunnel, recognizing specific sequence features and modulating translation speed. Our results challenge and extend the prevailing view of NAC’s function as a recruitment scaffold for cotranslational N-terminal acting factors such as SRP, METAP1,NatA, and NMTs to the ribosome^12,23,28–30^. Instead, NAC is a global chaperone that regulates elongation rate, assists domain-wise folding and promotes delivery of a subset of mitochondrial proteins and ER-destined proteins to their correct compartment. Remarkably, the pattern of NAC engagement is distinct for nascent chains destined for distinct cellular compartments.

**Figure 6.**
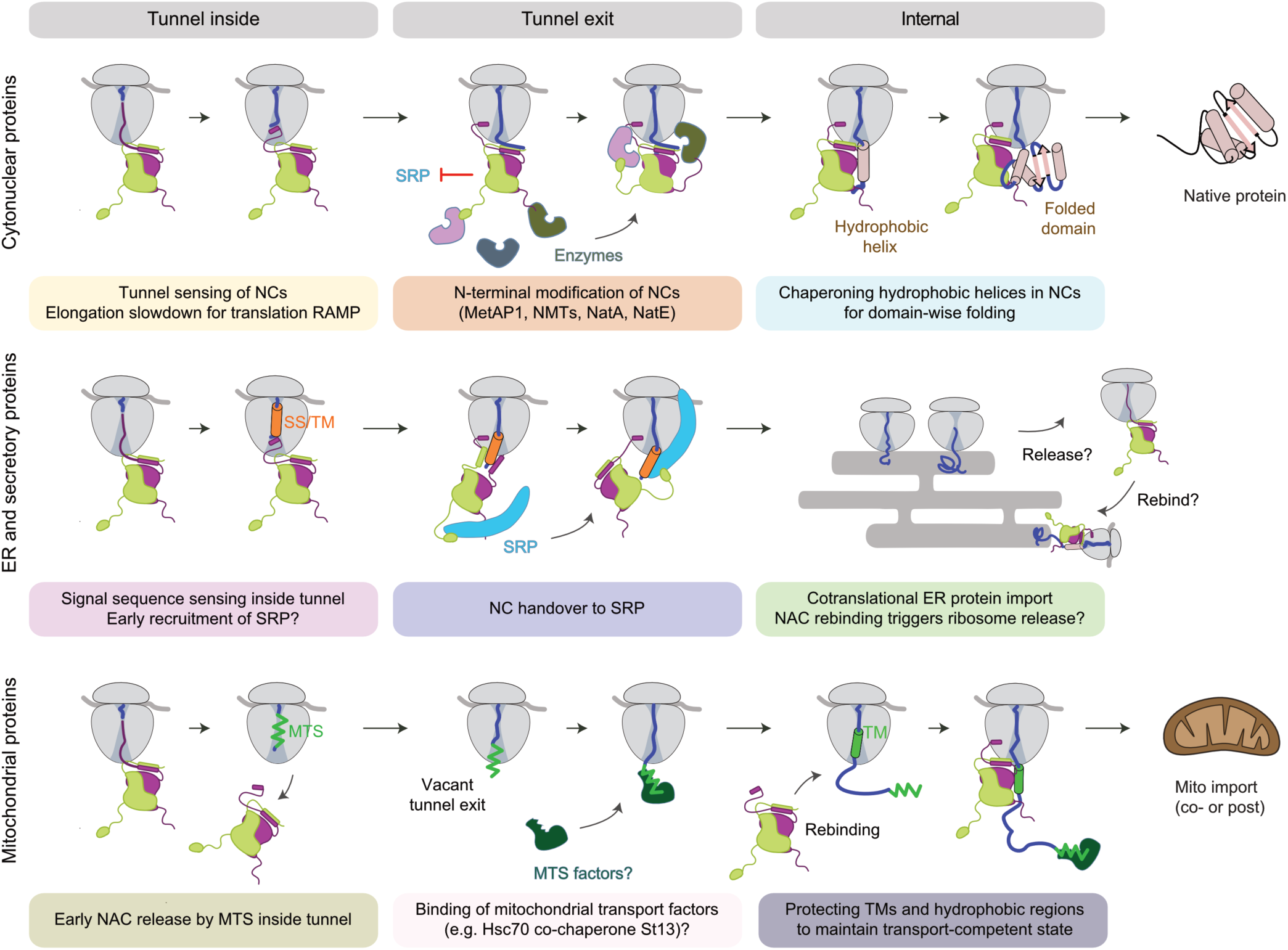
Model for NAC function during protein biogenesis. NAC acts in distinct manner depending on the localization of substrate. For cytonuclear proteins (top), NAC senses the nascent chains in the tunnel and induce elongation slowdown. Then recruits N-terminal modification enzymes for processing. As the nascent chain gets longer, NAC chaperones hydrophobic helices to assist domain-wise folding. For ER and secretory proteins (middle), NAC senses nascent chains in the tunnel and also induce elongation slowdown. As NC emerges out of the exit tunnel, NAC recruits SRP and relays the NC to SRP to be targeted to ER. At the ER, NAC may rebind to chaperone cytosolic segments during import. For mitochondrial proteins (bottom), NAC is absent in the beginning of translation. This may be caused by MTS or MTS binding factors that exclude NAC from ribosomes. As mitochondrial TMs emerge from the exit tunnel, NAC is recruited to chaperone hydrophobic TMs during targeting and import.

Our results define NAC as a chaperone that preferentially binds hydrophobic helices as they emerge from the ribosome. This recognition mode is distinct from that of Hsp70, which cotranslationally associates with hydrophobic elements with beta-sheets propensity^2,36^, as well as from TRiC, which binds cotranslationally to topologically complex domains^2^. The nonredundant specificity likely contributes to a tiered chaperoning system during translation. Because the exit tunnel environment favors helix formation^57^, NAC may engage nascent chains with hydrophobic content, regardless of their ultimate fold, priming them for subsequent folding or targeting. However, for cytonuclear proteins, this NAC recognition specificity appears to stabilize amphipathic and hydrophobic helices emerging from the ribosome to facilitate folding of specific domain classes (Fig. 6, top tier).. Further investigation into the interplay between different chaperones would shed light on how they cooperate to provide an optimal folding environment for nascent chains on the ribosome. The discovery of a universal NAC recognition motif raises interesting questions on which part of NAC is responsible for binding to the nascent chains and how NAC is able to sort them depending on their fate.

NAC is essential for biogenesis of a subset of mitochondrial protein. It does not interact with the MTS but instead engages hydrophobic helices and cofactor-binding motifs to maintain their solubility during cytosolic transit. This implies the existence of an unidentified mechanism, e.g. an MTS-binding factor, that prevent NAC binding, highlighting an unexplored layer of coordination in mitochondrial protein delivery. Our results also point to a more selective role for NAC in ER targeting. NAC does not interact with all ER-destined proteins, but instead binds to and enhances membrane delivery of specific ER-destined proteins through engagement with SS/TMs. While NAC and SRP have been shown to coexist on the ribosome^22,23^, our data shows that they engage SS and TM helices sequentially as they come out of the ribosome exit tunnel. Additionally, NAC’s ribosome association alone is sufficient to reduce mistargeting of cytosolic proteins to the ER, indicating a surveillance role independent of direct nascent chain interaction. These dual activities suggest that NAC plays both promotive and preventative roles in ER targeting depending on nascent chain identity.

We find that, unlike other chaperones such as Hsp70 and TRiC in eukaryotes or Trigger Factor in bacteria^2,36,58^, NAC also interacts with extremely short nascent chains still within the ribosome exit tunnel, consistent with previous in vitro structural studies of NAC^10^. These early tunnel interactions alter translation elongation dynamics, suggesting that NAC can tune ribosome flux through chaperone-mediated regulation. NAC slows elongation at early stages in a substrate-specific manner, correlating with amino acid sequence rather than codon usage. This reveals a new mechanism by which translation speed is modulated cotranslationally, broadening the known repertoire of elongation control beyond codon usage.

The novel regulatory function of NAC in translation elongation may serve several functions, including reducing ribosome density on some transcripts and regulating formation of disomes as well as other events which are tuned by elongation rate such as ER targeting. Our data indicates that NAC, rather than rare codons, is the primary driver of early elongation slowdown in *C. elegans*. We propose that NAC is responsible for the previously described “ramp” early in translation^31^. Our data supports emerging evidence that amino acid identity within the first five residues can modulate translation output^32^. NAC is the first known proteostasis factor that directly controls elongation rate, presumably through interaction with nascent chains. Our work opens the way for future structural studies with more fine-tuned nascent chain length and properties that will provide precise mechanisms of nascent chain sensing and discrimination by NAC both inside and outside the ribosomal tunnel.

Together, these findings establish NAC as a multifaceted regulator of early translation events, with roles in organelle targeting, chaperoning, and elongation control. Its unique ability to bind hydrophobic helices and interact within the ribosome exit tunnel underpins these diverse functions. Our findings resonate with previous data indicating that NAC has extra-ribosomal chaperone functions and can interact with amyloidogenic proteins such as Huntingtin and Aβ^59^. Given that NAC was implicated in aging and development^59–61^, with its levels decline with aging^62,63^, its loss may contribute not only to impaired protein targeting and folding but also to altered translation kinetics. This connection warrants further investigation into NAC’s role in age-related translation dysregulation and proteostasis collapse^54,63^.

## Methods

### Strains and growth conditions

Wildtype Bristol N2 strain was obtained from Caenorhabditis Genetics center. Worms were cultured according to standard techniques with E. coli OP50 as food source at 20°C^68^. Transgenic strains were generated using standard microinjection protocols^69^. Endogenous NACα locus (*icd-2*) was tagged with a Twin-Strep-FusionRed tag^70^ at its N-terminus using Homology-directed repair (HDR) of double-strand breaks induced by Cas9^71^. The spliced coding sequence of *slc-25a10* was amplified via PCR from N2 cDNA, C-terminally tagged with GFP and cloned into miniMos pCFJ910 vector^72^ under the control of bodywall muscle specific *unc-54* promoter and 3’utr. DNA was injected into worms at 20 ng/µl in the presence *myo2p*::mCherry (2.5 ng/µl) and DNA ladder (77.5 ng/µl, GeneRuler 1kb, Thermo Scientific). A single worm containing a stably inherited extrachromosomal array was selected for further analysis.

### RNAi

RNAi constructs targeting endogenous genes (NACα: *icd-2*; NACβ: *icd-1*, PAS-7: *pas-7*, TOMM-40: *tomm-40*) were cloned by inserting the spliced codon sequences into the vector L4440 (Addgene plasmid #1654). In double RNAi constructs, coding sequences were placed in tandem on a single plasmid. Constructs were transformed into the RNAi-feeding *E. coli* strain HT115(DE3) containing a modified lac operon for IPTG-induced expression of dsRNA^71^. Bacteria were induced by treatment with 1 mM IPTG for 2 h at 30°C before feeding to worms. Worms in liquid culture were fed fresh RNAi bacteria every day. For microscopy analysis, RNAi was performed from hatch on agar plates containing the respective induced HT115(DE3) RNAi bacteria. Nematodes were maintained on RNAi plates until the end of the experiment.

### Cloning and protein purification

Constructs for NAC expression and purification were described previously^10^. Briefly, wildtype and mutant human His-SUMO-NAC constructs were transformed into Rosetta (DE3) (Novagen). Cells were grown at 37C to OD_600_=1.5, cooled to 25C, and expression was induced with 1mM IPTG for five hours. Harvested cells were resuspended in lysis buffer (50mM sodium phosphate (pH 8), 300mM NaCl, 6mM MgCl_2_, 2mM β-mercaptoethanol, 2mM Pefabloc, 8 mg/ml Pepstatin A, 10 mg/ml Aprotinin, 5 mg/ml Leupeptin, 10 mg/ml DNase I, 10% glycerol), and lysed by high-pressure homogenizer (EmulsifFlex-C3). Cleared lysate was purified with Ni-IDA resin (Protino, Macherey-Nagel), incubated with 8μg Ulp-1 per mg protein to remove His-SUMO tag, followed by ion-exchange (ReSourceQ) to yield purified NAC complex with 1:1 ratio of αNAC and βNAC. p-benzoylphenylalanine (Bpa) incorporation into human NAC has been described previously^10^. Amber stop codons were inserted at the first position of NACβ of His-SUMO-NAC construct, co-transformed with Bpa/amber suppression-Rosetta plasmid into BL21(DE3)* (Novagen). Cells were grown at 37C to OD_600_=0.1, incubated with 1mM Bpa (Bachem) at 30C, and expression was induced at OD_600_ = 0.8 with 1mM IPTG for 12 hours at 20C. Proteins were purified from harvested cells following the same protocol as above.

### Microscopy

Worms were age synchronized and grown *eV/NACβ/tomm-40* RNAi until larval stage L4 before being shifted to plates co-knocking down the proteasomal subunit *pas-7.* On Day 3 of adulthood worms were immobilized on a 5 % agar pad with 10 mM levamisole. Worms were imaged using TCS SP8 MP (Leica) microscope with a HC PL APO CS2 63x/1.20 water objective. Images were processed in Fiji^73^. For mitochondrial stain, worms were fed overnight in the dark on fresh plates seeded with bacteria containing 1 µg/mL LumiTracker® Mito Red CMXRos (Lumiprobe). Worms were placed on fresh plates containing bacteria without LumiTracker® Mito Red CMXRos for 2h before microscopic analysis.

### Membrane fractionation

Lysis buffer (20 mM Tris-HCl, pH 7.4, 140 mM potassium chloride, 5 mM magnesium chloride, 1 mM DTT, 100 μg/mL cycloheximide, 20 U/mL SuperaseIn (Ambion) supplemented with Complete Protease Inhibitor Cocktail, EDTA-free (Roche)) was frozen dropwise in liquid nitrogen. One 4 mL aliquot of frozen lysis buffer was combined with 2 mL pellet of frozen worms in a 50 mL ball mill chamber (MM-301 CryoMill - Retsch) chilled in liquid nitrogen, using 2 rounds of 1 minute at 30 Hz. The pulverized worm material was divided into six equal portions for subsequent analyses (ribosome profiling and RNAseq). Pulverized worms were thawed in a room temperature water bath, supplemented with triton X-100 to 1% and incubated for 10 min on ice for lysis of the whole cell extract samples. Lysates were immediately centrifuged for 10 minutes at 20,000g, and the supernatant, constituting the whole cell extract, was collected. For the extraction of the total soluble fraction, additional pulverized worm samples were similarly thawed and immediately centrifuged at 20,000 xg for 10 minutes. The supernatant was collected as the total soluble fraction. The pellet remaining after centrifugation was resuspended in the lysis buffer supplemented with Triton X-100 to a final concentration of 1% and incubated for 10 min on ice and centrifuged under the same conditions as previously. The supernatant from this step, containing detergent-extracted membrane fractions, was retained. Total RNA from obtained whole cell, soluble, and membrane fractions were extracted using TRIzol reagent following manufacturer protocol.

### Immunoblot analysis and antibodies

For membrane fractionation samples, protein concentrations were normalized using the Bradford Assay (Bio-Rad), 4× NuPage LDS sample buffer (Thermo Fisher Scientific) was added, and samples were then boiled for 5 minutes. For gel electrophoresis, 10 ug of protein per sample was separated on a 4– 20% SDS–PAGE gel and transferred to a nitrocellulose membrane according to standard protocols. For NAC-bound ribosomes samples, 5X Lammli sample buffer was used and separated on 10% Bis-Tris gel. The membranes were blocked in 5% milk reconstituted in TBS (20 mM Tris pH 7.5, 150 ml NaCl, 0.1% NaN3). Primary antibodies were diluted in Antibody Buffer (20 mM Tris pH 7.5, 150 ml NaCl, 0.1% NaN3, 5% BSA) and secondary antibodies were prepared in 5% milk in TBS. As indicated, the blots were probed with specific primary antibodies against mouse Hsp-60 (DSHB), Pas-7 (DSHB, CePAS7), Actin (DSHB, JLA20), Sec61α (Santa Cruz, sc-393182), and Tubulin (DSHB, AA4.3, 1:20 dilution) and visualized using the LI-COR system IRDye 800CW donkey anti-mouse IgG (LI-COR 926-32212, 1:10,000 dilution). Commercial uL16 antibody was purchased from Biomol (#AP17603). *C. elegans* antiserum against NACαβ was raised in rabbits in-house.

### In vitro translation in Rabbit Reticulocyte Lysate and Bpa crosslinking

Non-stop mRNAs were generated by PCR as previously described^10^. Reverse primer encoded five C-terminal methionines to get robust radioactive signal from nascent chains with incorporated S^35^-Met, and terminal valine to stabilize tRNA-nascent chain^74^. Generated mRNAs were in vitro translated using 35 µl of rabbit reticulocyte lysate (Green Hectares) with 0.5-3µg of mRNA and S^35^-Met. RNCs were purified through 25% sucrose cushion in RNC wash buffer (50mM HEPES pH7.5, 500mM KOAc, 5mM Mg(OAc)_2_). Ribosomal pellet was resuspended in resuspension buffer (20mM HEPES pH7.5, 100mM KCl, 5mM MgCl_2_) and incubated with 10 µM Bpa-NAC variant at room temperature for 1 hour. Samples were crosslinked by UV irradiation for 30 minutes on ice. Crosslinked products were separated on SDS-PAGE and imaged using autoradiography.

### In vitro translation in reconstituted eukaryotic system

40S and 60S ribosomal subunits, all multisubunit eIFs, eEFs, and total aminoacyl-tRNA were purified from HeLa cells^55,75–77^. Single subunit eukaryotic initiation factors (eIF1, eIF1A, eIF4A, eIF4B, eIF5, eIF5A) were overexpressed in *E. coli* and purified according to published protocols^55,76,78^. Initiator Met-tRNA was fluorescence labeled by BODIPY-Fl as described^79^ . The first 150 nucleotides of the gene Y54E10BL.2.1 were cloned in pUC19 under the proper Kozak context with an unstructured 5’ end to allow efficient translation with the minimal set of translation factors^55,80^. mRNA was *in vitro* transcribed with T7 RNA-polymerase and purified by cation exchange chromatography followed by ethanol precipitation. Initiation complexes (IC) were assembled in buffer containing 20 mM HEPES, pH 7.5, 95 mM KOAc, 3.75 mM Mg(OAc)_2_, 1 mM ATP, 0.5 mM GTP, 0.25 mM spermidine, 2 mM DTT. 48S IC was prepared by incubating 40S (0.22 µM) with eIF1 (1 µM), eIF1A (1 µM), eIF2 (0.75 µM), eIF3/eIF4E/eIF4G (0.75 µM eIF3, 0.3 µM eIF4G, 0.23 µM eIF4E, i.e., keeping the major eIF4F components in about 1:1 ratio to the 40S subunit), eIF4A (1 µM), eIF4B (0.44 µM), BODIPY-Met-tRNA (0.22 µM), and mRNA (0.66 µM) for 10 min at 37°C. To form 80S IC, eIF5 (5 µM), eIF5B (1 µM) and 60S (1 µM) were added and incubated for 5 min at 37°C. For ternary complex (TC) formation, eEF1A (25-55 µM) was incubated with GTP (1 mM), creatine phosphate (3 mM) and creatine phosphokinase (0.5 mg/mL) and total aminoacyl-tRNA (1.7-2.7 OD_260_) for 15 min at 37°C. Elongation factor mix consisting of TC, eEF2 (1 µM) and eIF5A (2 µM) was mixed with 80S IC in the translation buffer (20 mM HEPES pH 7.5, 2.5 mM Mg(OAc)_2_, 120 mM KOAc, 0.25 mM spermidine, 2 mM DTT) at 37°C to start translation elongation. The translation was done in the absence (compensated with buffer) or in the presence of 8 µM WT or mutant NAC. The incubation was carried out for 60 s with an aliquot taken every 2 s to follow peptide chain elongation in real time. The reaction was quenched by 0.33 M NaOH and incubated for 30 min to hydrolyze all nascent peptides, followed by pH neutralization with a final concentration of 0.31 M HEPES. The N-terminal BODIPY FL-labeled translation products were analyzed using three-layered Tris-Tricine-SDS PAGE^81,82^ and visualized at 488 nm using Typhoon scanner. The bands were quantified using Image Studio, normalized against a loading control (fluorescently labeled *E. coli* ribosomal protein S6), and analyzed in GraphPad prism.

### RNA-Seq library preparation and RNA-Seq sequencing

Total RNA integrity and concentration assessed using the Agilent Bioanalyzer 2100, employing the RNA 6000 Nano Kit (Agilent Technologies, Cat. No. 5067-1511). Ribosomal RNA was depleted from 1 µg total RNA using Illumina’s Ribo-Zero rRNA removal Kit (Document # 15066012 V02). Library prep was performed with Illumina’s the TruSeq Stranded Total RNA Prep Kit, following the manufacturer’s instructions (Document # 1000000092426v01) and starting with the Elute, Fragment, Prime, High Conc Mix, and fragment distribution was assessed using the DNA 1000 chip on the Agilent Technologies 2100 Bioanalyzer. Libraries were amplified, profiled with a High Sensitivity DNA Chip on a 2100 Bioanalyzer (Agilent Technologies), and quantified by qPCR (KAPA Library Quantification Kits for Illumina platforms v1.14). Samples were pooled in equimolar ratio and sequenced on a NextSeq 550 (Illumina).

### Isolation of NAC-bound ribosomes

Worms expressing Twin-Strep tagged NACα were age-synchronized by 20% alkaline hypochlorite treatment, grown in liquid culture for 44 h at 20°C and harvested on ice in 0.1 M NaCl. Separation from *E. coli* food was achieved by sucrose flotation and a subsequent wash in 0.1 M NaCl. Worms were extracted in the presence of 10 mM DSP (Pierce) in RNC buffer (30 mM HEPES pH 7.4, 100 mM KOAc, 5 mM MgCl_2_, 5% Mannitol, 100 µg/ml Cycloheximide, 1x Protease inhibitor cocktail (Roche)) by mild sonication on ice. Lysate was cleared by centrifugation (10 min, 12000 g, 4°C), filtration of the supernatant through a 0.45 µm nitrocellulose membrane and centrifuged again (20000 g, 5 min). Ribosome nascent chain complexes (RNC) were sedimented through a 30 % sucrose cushion prepared in RNC buffer by ultracentrifugation for 3.5 h at 60000 rpm in a 70.1 Ti rotor (Beckman). RNCs were resuspended in RNC buffer. After quantifying RNA concentration, mRNA was digested with RNase I (Ambion) for 45 min at RT using 0.4 U/µg RNA. RNase reaction was stopped by addition of Superase•In (Invitrogen) at 0.4 U/µg RNA. Samples of total RNA were removed for ribosome profiling of the total translatome. The NAC-bound fraction of RNCs was recovered via affinity purification over a 250 µl column of Strep-Tactin (IBA, Germany) and thoroughly washed with RNC buffer containing 0.02 U/µl Superase•In. Elution was achieved by addition of 20 mM Desthiobiotin (IBA, Germany) in RNC buffer containing 0.02 U/µl Superase•In. Total and NAC-bound RNCs were loaded on a 15 -45 % sucrose gradient prepared in RNC buffer containing 0.02 U/µl Superase•In. Gradients were subjected to ultracentrifugation for 15 h at 16000 rpm on a SW 41 Ti rotor (Beckman) and fractionated from top to bottom with a density gradient fractionator (Brandel). Fractions containing monosomes were collected and ultracentrifuged (1.5 h, 200000 g) in a S140-AT rotor (Sorvall). Digested NAC-bound RNCs and total RNCs were resuspended in RNC buffer containing 0.02 U/µl Superase•In and flash frozen in liquid nitrogen.

### Ribosome profiling sample preparation

Ribosome profiling libraries were prepared following previously published protocol with modifications^2,52^. For analyzing changes in translation kinetics upon NAC-KD, worms at L4 stage were rapidly collected and frozen. Frozen cells were lysed using Cryo-Mill (Retsch, MM301) in the presence of 2ml of lysis buffer (20 mM Tris-HCl pH 7.5, 140 mM KCl, 5 mM MgCl2, 1 mM DTT, 100 µg/ml Cycloheximide, 1% Triton X-100, and 1 X Protease Inhibitor). Lysed powder was quickly thawed in a water bath at room temperature and spun at 21,000 g for 15 minutes at 4 °C to clear lysate. RNAse I (Invitrogen, AM2294) was added to 0.4U/μg of RNA and incubated at 25 °C for 45 minutes. Digestion was stopped by adding 0.4U/μg of SUPERaseIn RNAse Inhibitor (Invitrogen, AM2696). RNAse-treated lysate was layered on 900 μl sucrose cushion buffer (20 mM Tris-HCl pH 7.5, 140 mM KCl, 5 mM MgCl2, 1 mM DTT, 100 µg/ml Cycloheximide, 0.02U/μl SuperaseIn, 1M Sucrose), and spun at 100,000 rpm for 1 hour at 4 °C in TLA100.3 rotor. Resulting ribosome pellet was resuspended in 250 μl of lysis buffer with SuperaseIn and RNA was extracted using TRIzol reagent (Invitrogen, 15596026) following manufacturer’s protocol. Similarly, for selective ribosome profiling, RNA was extracted from isolated NAC-bound ribosomes. 27-34bp fragments were isolated from denaturing gel, ligated to adapter (NEB, S1315S), and ribosomal RNA was removed using Ribo-Zero (illumina) mixed with custom depletion DNA oligos (Table 1). Remaining fragments were reverse transcribed, circularized, and PCR amplified following the steps described previously^52^. Barcoded samples were pooled and sequenced using Nextseq 550 (Illumina).

**Table 1:**
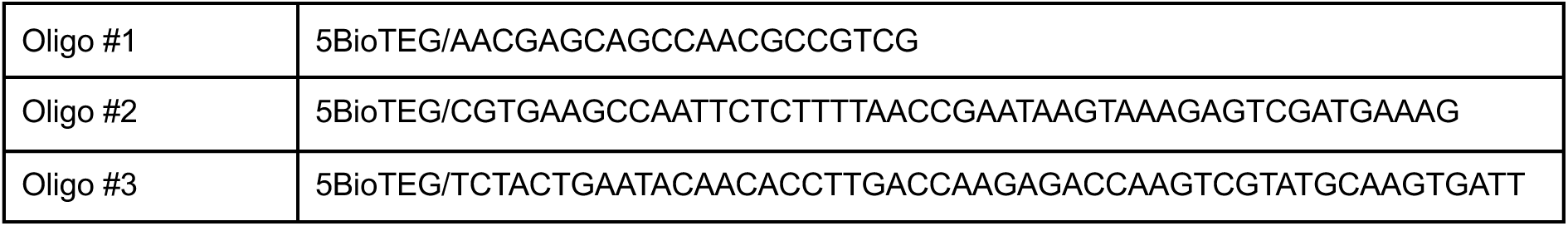
List of DNA oligonucleotides used for ribosomal RNA depletion.

### Data Analysis

#### Data processing

Data processing was done following previously published protocol with minor modifications(Stein, Kriel and Frydman 2019). Adapter sequences were removed from demultiplexed sequencing reads using Cutadapt v.1.4.2(Martin 2011), followed by the removal of the 5’ nucleotide using FASTX-Trimmer. Reads mapping to ribosomal RNAs were removed using Bowtie v.1.3.1(Langmead, Trapnell et al. 2009). Remaining reads were aligned to the reference transcript library that consisted of coding sequences containing 21 nucleotides flanking upstream of the start codon and downstream of the stop codon. To maximize unique mapping, a reference transcript library was constructed using the longest isoform for every 20145 gene in the WBcel235 genome assembly. Bowtie alignment was performed using the following parameters: -y -a -m 1 -v 2 -norc - best -strata. A-site offset was estimated using riboWaltz(Lauria, Tebaldi et al. 2018), and fragment lengths that do not exhibit 3-nucleotide periodicity were removed. Pause scores at each position were calculated by dividing the number of reads at each position by the average number of reads within the internal part of the transcript, excluding the first and last five codons.

#### NAC binding site identification

NAC binding sites were identified following established protocol^2^. To increase confidence on positional enrichment of NAC, only transcripts with good correlation between replicates (Pearson’s r ≥ 0.5 based on RPM) and good coverage (≥0.5 reads per codon and ≥64 reads per gene) were analyzed (NAC_WT_: 4425 gene, NAC_ΔN1-53_: 2128 genes). Replicates of resulting transcripts were summed to increase coverage. Using raw read counts from total and pulldown samples, we created 2 x 2 contingency tables at each position of a given transcript, and performed two-tailed Fisher’s exact tests to compare the ratio of reads at each position to the ratio at all other positions of a given transcript (odds ratio). The resulting p values were adjusted using Benjamini-Hochberg correction for multiple hypothesis testing per transcript. We used established thresholds^2^ to identify positions of significant NAC enrichment (1 < odds ratio < ∞, padj < 0.05, and pulldown reads > average reads across transcript to control for the background). Finally, NAC binding events were identified as positions that pass the thresholds for at least five consecutive codons, except for those occurring within the first 30 codons of a transcript whose threshold was lowered to three consecutive codons. Positions with the highest enrichment (odds ratio) within a binding event (consecutive codons) were identified to yield 2578 unique binding events within 1399 transcripts for NAC_WT_ and 2265 unique binding events in 1057 transcripts for NAC_ΔN1-53_.

#### NAC enrichment analysis

NAC enrichment of individual transcripts was plotted by first smoothing the raw reads in 15 codon window to avoid division by 0, calculating odds ratio, and plotting moving average of odds ratio in a window of 15 codons to reduce noise. Note that some enrichment peaks appearing in these plots are not statistically significant. For average enrichment plots, average odds ratio of identified substrates around respective motifs was plotted with bootstrapped 95% confidence interval.

#### Domain Analysis

Domain annotations for *C. elegans* were downloaded from PFAM^1^ and CATH/Gene3D^3,37^. Identified NAC binding sites were mapped to domain boundaries, and assigned to respective domain if binding sites falls between the start and 30 codons after the end of the domain to account for the portion covered by ribosome exit tunnel. Enrichment and p values were calculated by comparing occurrence of specific domain within the transcripts analyzed and occurrence of specific domain within assigned NAC binding sites.

#### Physiochemical properties of nascent chain

Positional net charge was calculated as average charge within six codon window following Lehninger scale using Peptides package^83^. Positional hydrophobicity was calculated as average hydrophobicity at a given position following Kyte-Doolittle scale using Peptides package, and plotted as moving average in a window of six codons. Hydrophobicity and bulkiness of signal sequences and transmembrane helices were calculated using Peptides package. To visualize amino acids enriched around NAC binding sites, we used the weighted Kullback Leibler method^84^ using the frequency of each amino acid in coding sequences of reference as a background.

#### Identification of transcripts with NAC-dependent slowdowns

For each transcript, 2 x 2 contingency table was generated by summing raw reads within or outside of respective regions (Pre-constriction: 1-10 or Post-constriction: 11-30) in each condition (WT or NAC-KD). Two-tailed Fisher’s exact test was performed to calculate the odds ratio and p value. P-value was adjusted using Benjamini-Hochberg correction for multiple hypothesis testing. Replicates were analyzed separately and only transcripts that have odds ratio <1 and padj < 0.05 in both replicates were selected as having NAC-dependent slowdowns.

#### Membrane fractionation and RNA-seq

Demultiplexed and adapter-removed sequencing data was aligned to the same transcript reference described above using STAR^85^. Resulting counts data for total lysate were used for calculating translation efficiency. Resulting counts data for soluble and membrane from each replicate were used to calculate the fraction membrane bound, and the average value of two replicates was used. The targeting defect was quantified by the ratio of the fraction membrane bound between the two conditions (NAC-KD/WT).

#### Miscellaneous annotations and software

Secondary structure was predicted by running PSIPRED^86^ on the whole protein sequence. Subcellular localization annotation was downloaded from LocTree^87^. Signal sequence and transmembrane domain annotations were directly downloaded from Uniprot website. Common predictions of MTS by MitoFates^5^ and TargetP^6^ were used.

#### Cross-validation of NAC binding site identification

We performed a parallel analysis using a different method^67^ to ensure that the conclusions drawn from analyzing SeRP data is not dependent on the method of analysis. Data was processed and analyzed as described in ^67^. Briefly, after removing the adapters and non-coding RNAs as described above, reads were aligned to genome using STAR^85^. P-site offset was applied using custom Julila script provided in ^67^. NAC binding positions were identified based on the threshold (1.5) method. Method based on background model was not used because NAC does not show depletion in the first 30 amino acids. The conclusions from parallel analysis are shown in Extended Data Fig. 9.

## Author contributions

J.H.L., J.F., L.R., M.G., E.D., and M.V.R. designed the study. J.H.L., L.R., M.G., and V.G. performed all of the experiments and computational analyses, with assistance from K.M.K., J.A., and A.K. in carrying out crosslinking experiments, N.M.B. and F.M.P. in carrying out Ribosome profiling and RNA-seq, and E.S. in carrying out reconstituted eukaryotic in vitro translation experiments. J.H.L. and J.F. wrote the manuscript with input from all authors.

## Acknowledgements

We thank Olaf Geintzer, Vanessa Herold, Sandra Kappler, Christina Kothe, Anna Pfeifer, Theresia Steiger, Franziska Hummel and Michael Zimmermann for expert technical assistance. We thank Dr. K. Annamalai for providing the C. elegans MTS-CFP strain. This research was supported by grants from NIH (R01GM056433) and NIA (P01AG054407) to J.F., NIA (T32AG000266) and startup funds from Stony Brook University to J.H.L., the German Science Foundation (DFG): SFB969/A01/A07 and 537004599 to E.D. and M.G, Priority Program (SPP2453) 541647156 to E.D., the Max Planck Society and a grant of the Deutsche (SFB1565, P15, project number 469281184) and the Leibniz Prize of the Deutsche Forschungsgemeinschaft to M.V.R.

---

**Extended Data Figure 1.**
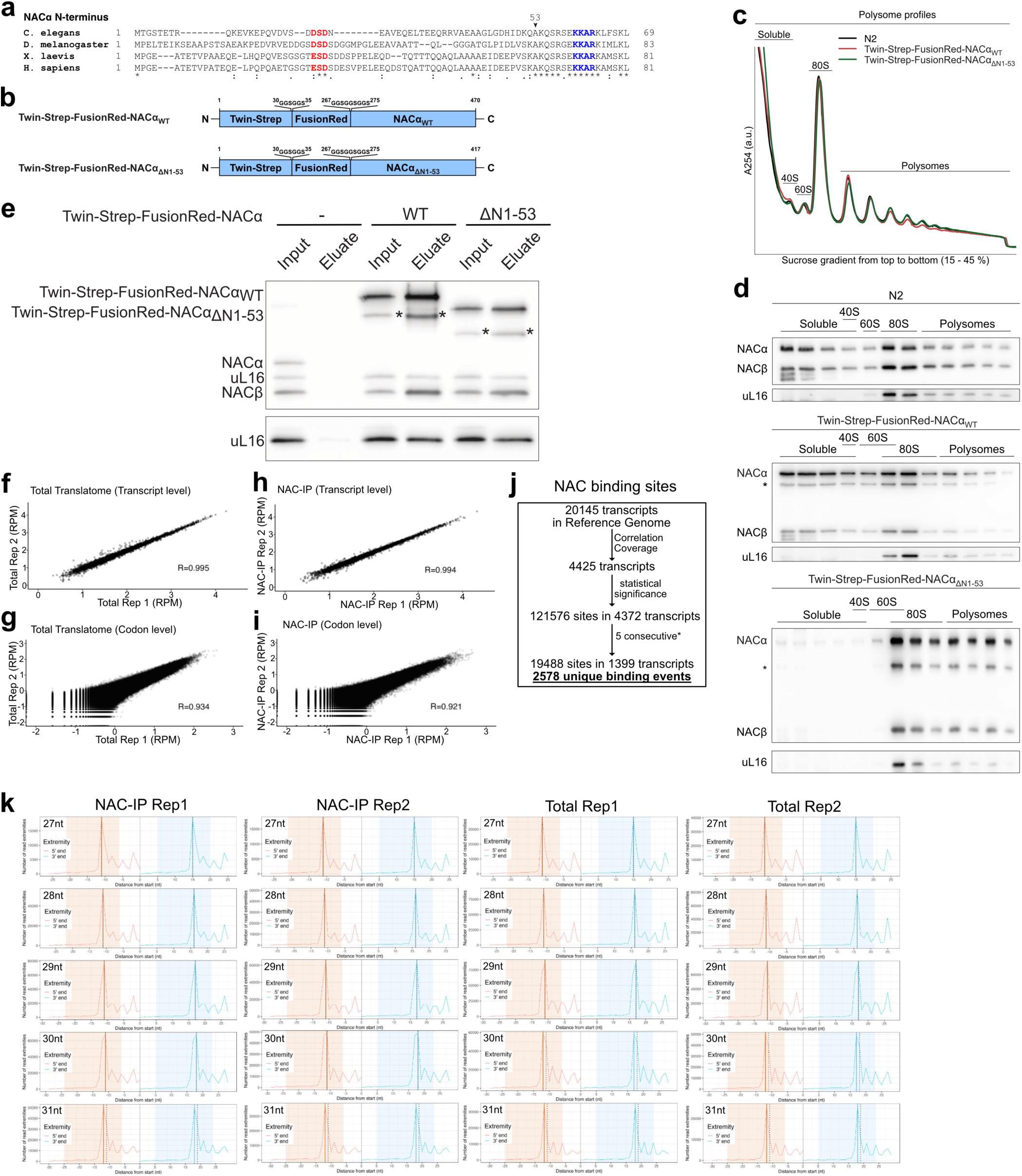
Pulldown of NAC-bound ribosomes and binding site identification. **a,** Sequence conservation of NACα N-terminus showing the region that negatively regulates ribosome interaction through negatively charged patch^10^. **b,** NAC constructs used for isolating NAC-bound ribosomes. Twinstrep and FusionRed were separated with GS linker to ensure accessibility of the tag and minimize effect on NAC function. **c,** Sucrose gradient fractionation of ribosomes from worms expressing WT (N2, *blue*), Twin-Strep-FusionRed-NACα (*red*), or Twin-Strep-FusionRed-NACα_ΔN1-53_ (*green*). Profiles from all conditions are identical. **d,** Western blot to show distribution of wild-type or NAC variants across the sucrose fractionation profiles. Twin-Strep-FusionRed-NAC is distributed across the gradient similar to WT, while NACα_ΔN1-53_ is almost exclusively bound to ribosome. *Cleavage fragment of FusionRed described previously^10^. **e,** Input and elution of NAC-bound ribosomes from untagged (left), Twin-Strep-FusionRed-NACα (middle), or Twin-Strep-FusionRed-NACα_ΔN1-53_ (right). uL16 was detected in the top blot as it runs in between NACα and NACβ. *Cleavage fragment of FusionRed described previous^64^. **f-i,** Correlation between replicates of total translatome (**f**-**g**) or NAC-IP (**h**-**i**) at transcript (**f** and **h**) or codon (**g** and **i**) level. **j,** Schematic of NAC binding site identification. Out of 20145 unique transcripts, 4425 that have good coverage and reproducibility between replicates, were analyzed to compare total translatome and NAC-IP datasets and identify positions with statistically significant enrichment in NAC-IP dataset. These positions were further filtered to include sites that show at least five consecutive codons of significant enrichment to find 2578 binding events in 1399 transcripts. *Note: Positions with at least three consecutive codons of significant enrichment were used for binding sites within the first 30 residues. **k,** Output from riboWaltz package showing the periodicity and p-site estimation.

**Extended Data Figure 2.**
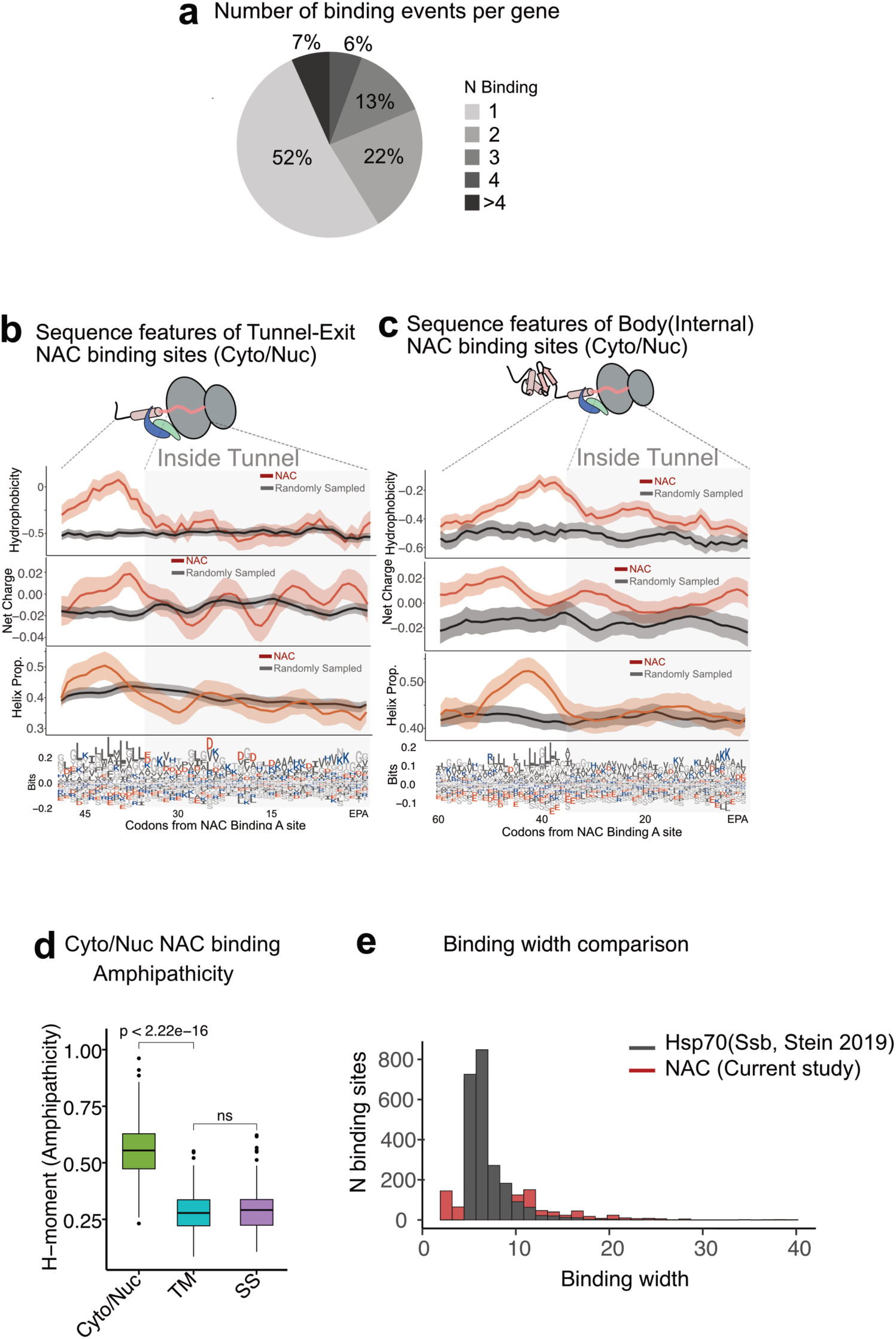
NAC interactions with cytosolic proteins. **a,** Number of NAC binding events per protein for cytonuclear NAC substrates. **b-c,** Average of physiochemical properties of nascent chains aligned to NAC binding sites in Tunnel-exit (**b**) or Internal (**c**) regions. The shaded region represents a region buried inside the ribosome exit tunnel that spans 35 amino acids of nascent chains. **d**, Hydrophobic moment (Amphipathicity) of NAC interacting regions of cytonuclear proteins (n=723) or ER targeting sequences (1^st^ transmembrane helices (TM, n=109) and signal sequences (SS, n=190)). Statistical analysis was performed using two-sided t-tests. **e,** Binding width of NAC in comparison to binding width of Hsp70 (Ssb) that was studied previously.

**Extended Data Figure 3.**
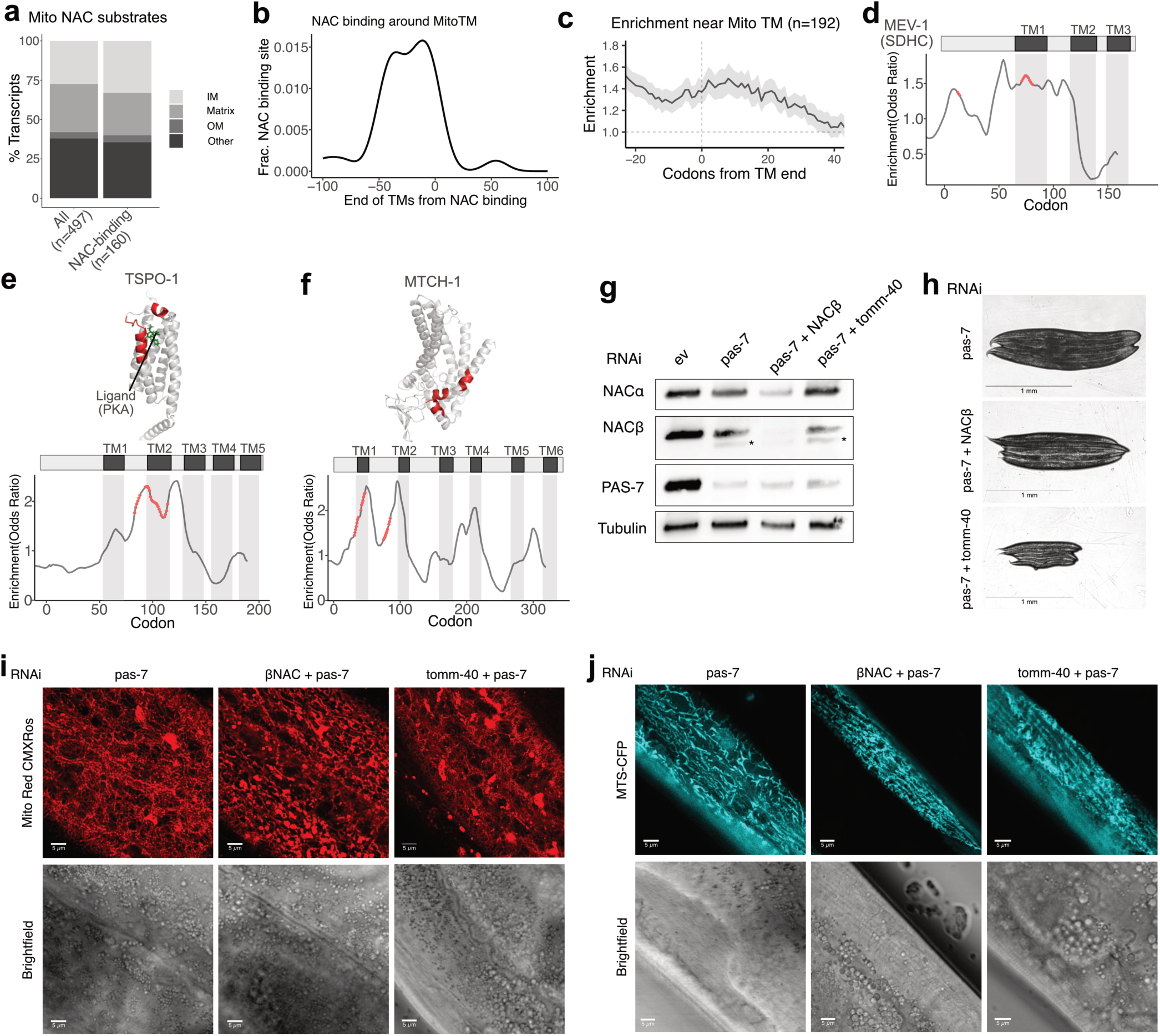
Role of NAC in mitochondrial protein biogenesis. **a,** Distribution of sub-organelle localization of mitochondrial proteins for all mitochondrial proteins or those that interact with NAC. IM: inner-membrane, OM: outer-membrane, Other: not annotated. **b**, Positions of NAC binding sites relative to the end of mitochondrial TMs. Fraction of NAC binding sites peaks just before 0, indicating NAC interacts as the TMs emerge out of the exit tunnel. **c,** Enrichment of NAC around mitochondrial transmembrane segments. **d-f,** Example mitochondrial membrane proteins MEV-1(SDHC) (**d**), TSPO-1^65^ (**e**), and MTCH-1 (**f**) that interact with NAC near transmembrane helices. Transmembrane helices are shaded, and NAC binding regions are highlighted in red in **e** and **f**. **g,** Western blots of NACα, NACβ, and PAS-7 to show the levels of proteins under different knockdown conditions. *Degradation product of NACβ that is stabilized upon pas-7 knockdown. **h,** Size of worms under different knockdown conditions. tomm-40 knockdown shows a strong phenotype suggesting high efficiency of knockdown. **i,** Microscopy visualizing mitochondria using MitoTracker Red CMXRos with knockdown of pas-7 (left), NACβ+pas-7 (middle), or tomm-40+pas-7 (right). **j,** Microscopy visualizing CFP (Cyan Fluorescent Protein) fused to mitochondrial targeting sequence (MTS) [myo-3p::MtLS::CFP] with knockdown of pas-7 (left), NACβ+pas-7 (middle), or tomm-40+pas-7 (right).

**Extended Data Figure 4.**
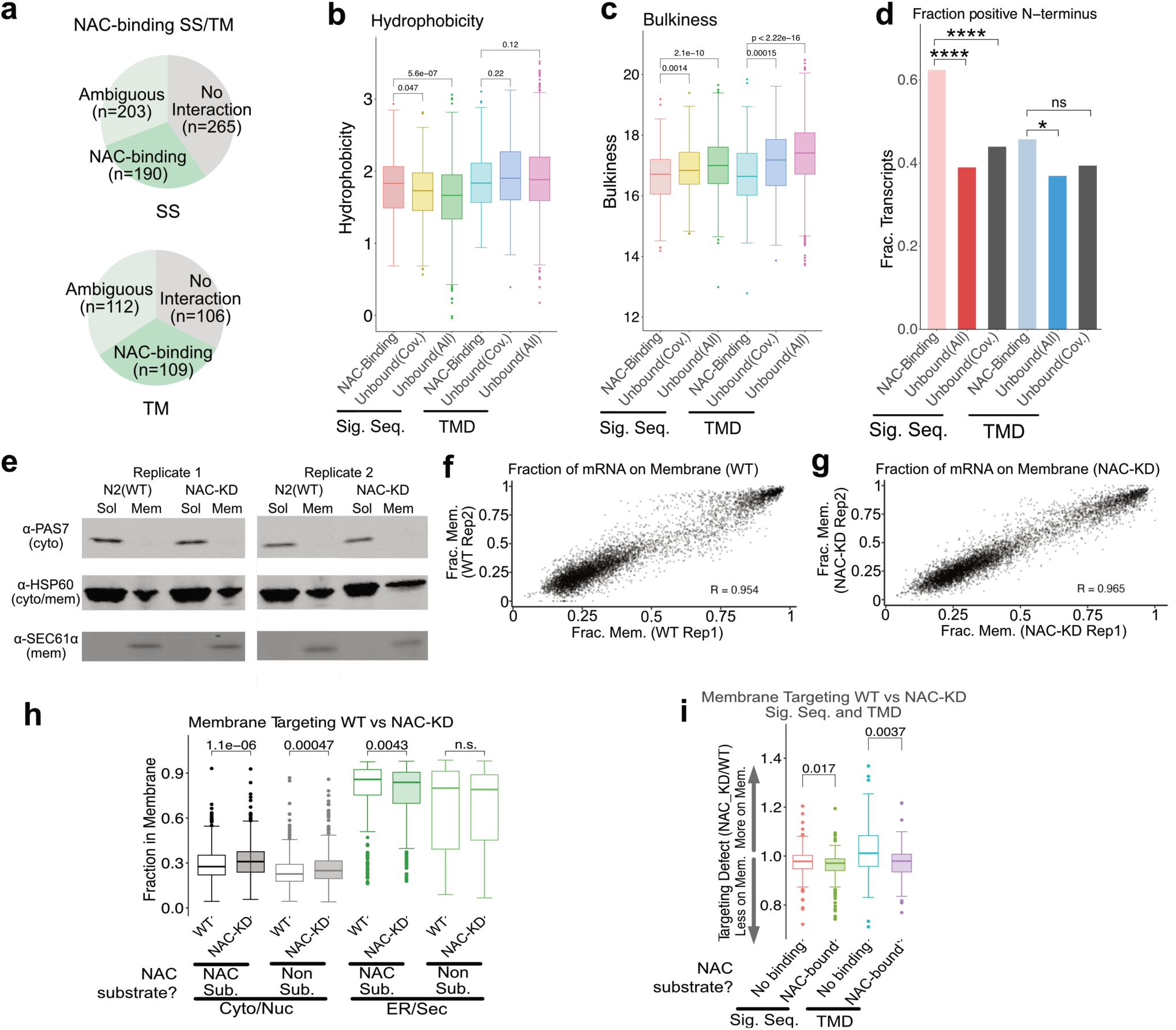
Role of NAC in cotranslational ER localization of nascent proteins. **a,** Fraction of ER-localized proteins with N-terminal transmembrane helices (TM) or signal sequence (SS) that interact with NAC during synthesis. Ambiguous group represents proteins that show some degree of NAC enrichment but not enough to pass the threshold (see Methods). **b-d,** Hydrophobicity (**b**), Bulkiness (**c**), or fraction of N-terminal positive charge (**d**) of signal sequences (SS) or transmembrane helices (TM) that interact with NAC (SS: n=190, TM: n=109) or those that do not (unbound) (SS: n=265 (Cov)/n=2809 (All), TM: n=106(Cov)/n=4063(All)). ‘(Cov)’ is among a subset of 4425 transcripts that had sufficient coverage which were analyzed, while ‘(All)’ is a subset of all 20145 unique transcripts in the reference transcriptome. Statistical analysis was performed using two-sided Wilcoxon rank-sum tests for hydrophobicity and bulkiness, and Fisher’s exact test for fraction positive N-terminus (Sig. Seq.:p=2.6x10^-9^(All), p=4.6x10^-6^(Cov), TM: p=0.022(All), p=0.89(Cov)). **e,** Western blot of soluble (Sol) and membrane (Mem) fraction isolated without (left) or with (right) NAC knockdown. PAS-7 was used as soluble, HSP-60 as both soluble and membrane, and SEC61α as a membrane marker. **f-g,** Scatter plot of RNA-seq from membrane fractionation of WT (**f**) or NAC-KD (**g**), showing the reproducibility of the data. **h,** Fraction membrane localized Cyto/Nuc (NAC Sub.: n=624, Non Sub.: n=442) or ER/Sec (NAC Sub.: n=366, Non Sub.: n=277) mRNA for NAC substrates and non-substrates under WT or NAC-KD condition. Statistical analysis was performed using two-sided Wilcoxon rank-sum tests. **i,** Defect in membrane targeting caused by NAC-KD for signal sequence (SS) (NAC-bound: n=175, No binding: n=142) or transmembrane helices (TM) (NAC-bound: n=101, No binding: n=74) containing proteins that do or do not interact with NAC. Statistical analysis was performed using two-sided Wilcoxon rank-sum tests.

**Extended Data Figure 5.**
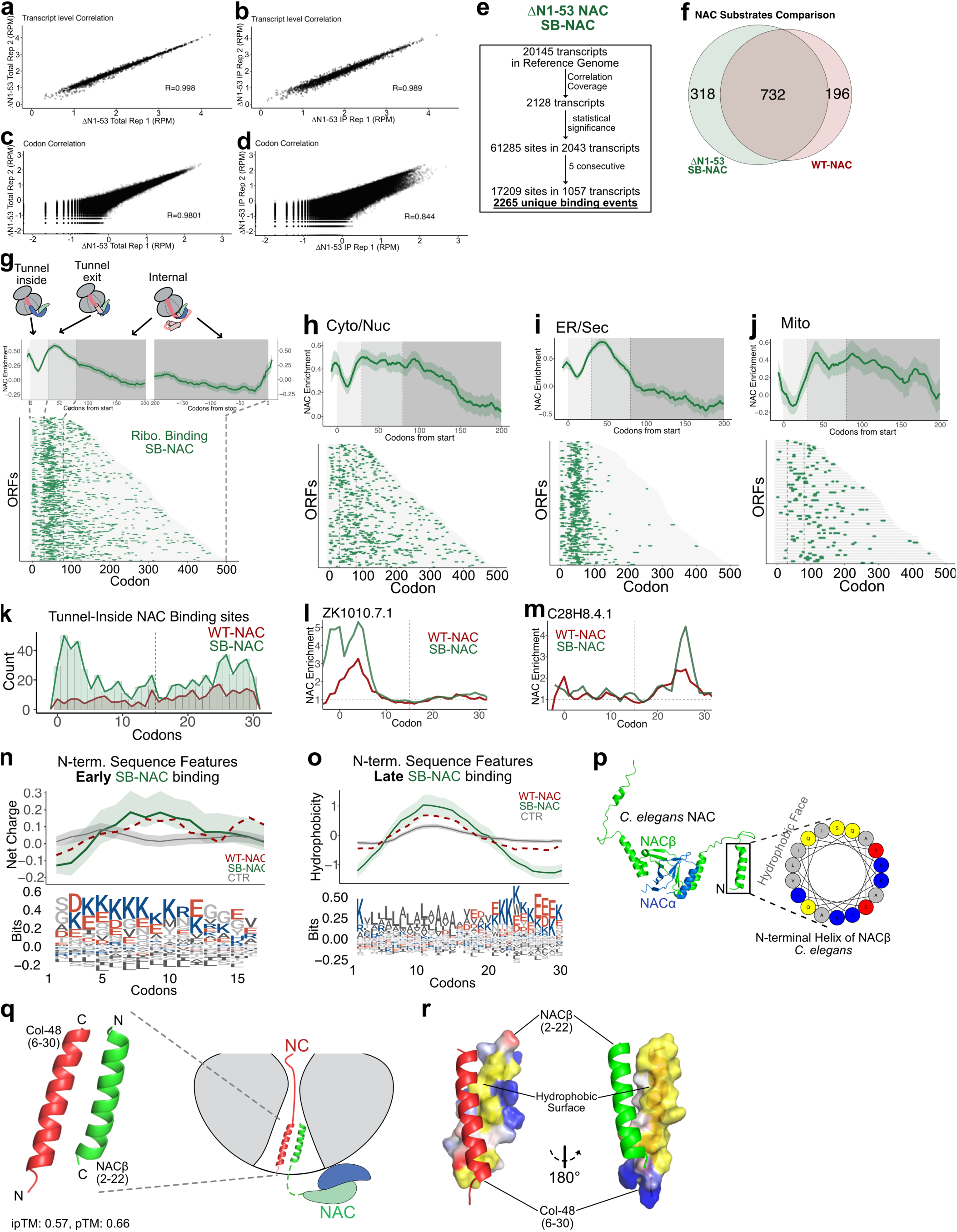
Characterization of mutant NAC_ΔN1-53_ interaction with translating ribosomes. **a-d,** Correlation between replicates of total translatome (**a** and **c**) or mutant NAC_ΔN1-53_-IP (**b** and **d**) at transcript (**a-b**) or codon (**c-d**) level. **e,** Schematic of NAC binding site identification. Out of 20145 unique transcripts, 2128 that have good coverage and reproducibility between replicates, were analyzed to compare total translatome and NAC-IP datasets and identify positions with statistically significant enrichment in NAC-IP dataset. These positions were further filtered to include sites that show at least five consecutive codons of significant enrichment to find 2265 binding events in 1057 transcripts. **f,** Overlap between substrates identified from wildtype NAC (*red*) and mutant NAC_ΔN1-53_ (*green*). Only transcripts analyzed in both datasets were included. **g-j,** Positions of mutant NAC_ΔN1-53_ interaction (*green*) with nascent proteome in three regions; Tunnel inside (codon <30), Tunnel exit (30 < codon < 80), and Internal (codon>80) for all (**g**), Cyto/Nuc (**h**), ER/Sec (**i**), Mito (**j**). **k,** Histogram of number of NAC binding sites identified within the first 30 codons from NAC_WT_ (*red*) or NAC_ΔN1-53_ (*green*). **l-m**, Enrichment of NACWT (red) or NACΔN1-53 (green) on example proteins ZK1010.7.1 (**l**) and C28H8.4.1 (**m**). **n,** Sequence features of N-termini of nascent chains that interact with mutant NAC_ΔN1-53_ during the synthesis of first 15 amino acids (**Early**, *green*) or randomly sampled (CTR, *grey*). Data from NAC_WT_ (*red*) is overlayed for comparison. **o,** Sequence features of N-termini of nascent chains that interact with mutant NAC_ΔN1-53_ during the synthesis between 15 and 30 amino acids (**Late**, *green*) or randomly sampled (CTR, *grey*). Data from NAC_WT_ (*red*) is overlayed for comparison. **p,** Structure of NAC^23^ and helical wheel diagram of N-terminal helix of NACβ showing amphipathic feature. Residues are colored according to their physical properties on helical wheel diagram (hydrophobic (*grey*), polar (*yellow*), basic (*blue*), acidic (*red*)). **q,** Model for interactions between N-terminus of a protein that interacts with NAC inside ribosomal tunnel (Collagen protein, Col-48, *red*) and N-terminus of NACβ (*green*) was predicted with AlphaFold with a good confidence ipTM:0.57 and pTM=0.66. This illustrates how NAC may interact with nascent chains towards the vestibule of the tunnel. **r,** electrostatic surface representation to highlight the hydrophobic surface (yellow) of NACβ and Col-48.

**Extended Data Figure 6.**
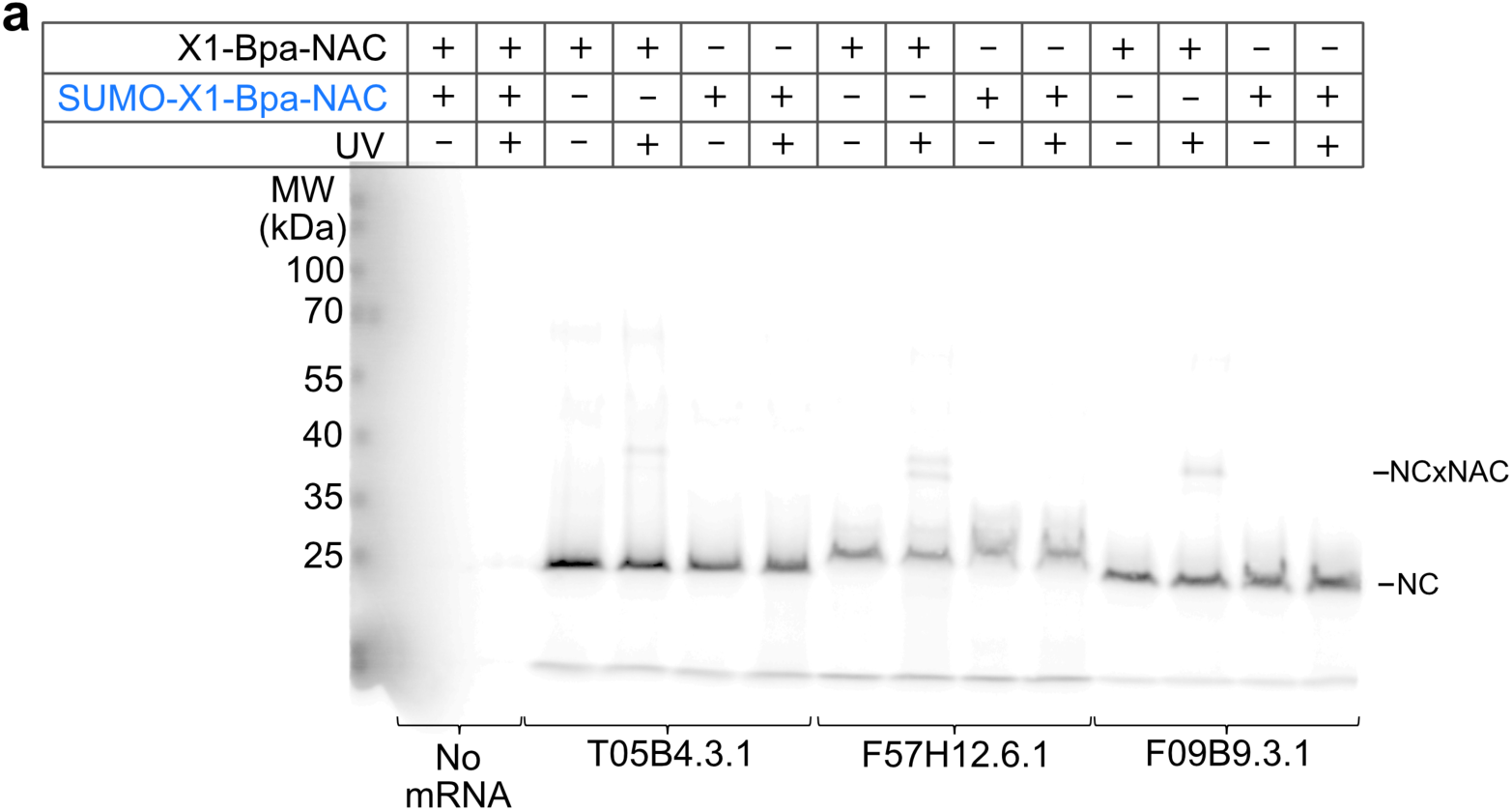
Bpa-crosslinking of NAC and ribosome nascent chain complexes. **a,** Experimental setup is identical to as in Fig. 4g except a NAC variant with SUMO domain fused to N-terminus of NACβ was used to prevent insertion into the ribosome exit tunnel. Uncrosslinked nascent chain (NC) migrates ∼25kDa while NC crosslinked to NAC (NCxNAC) migrates just below 40kDa marker.

**Extended Data Figure 7.**
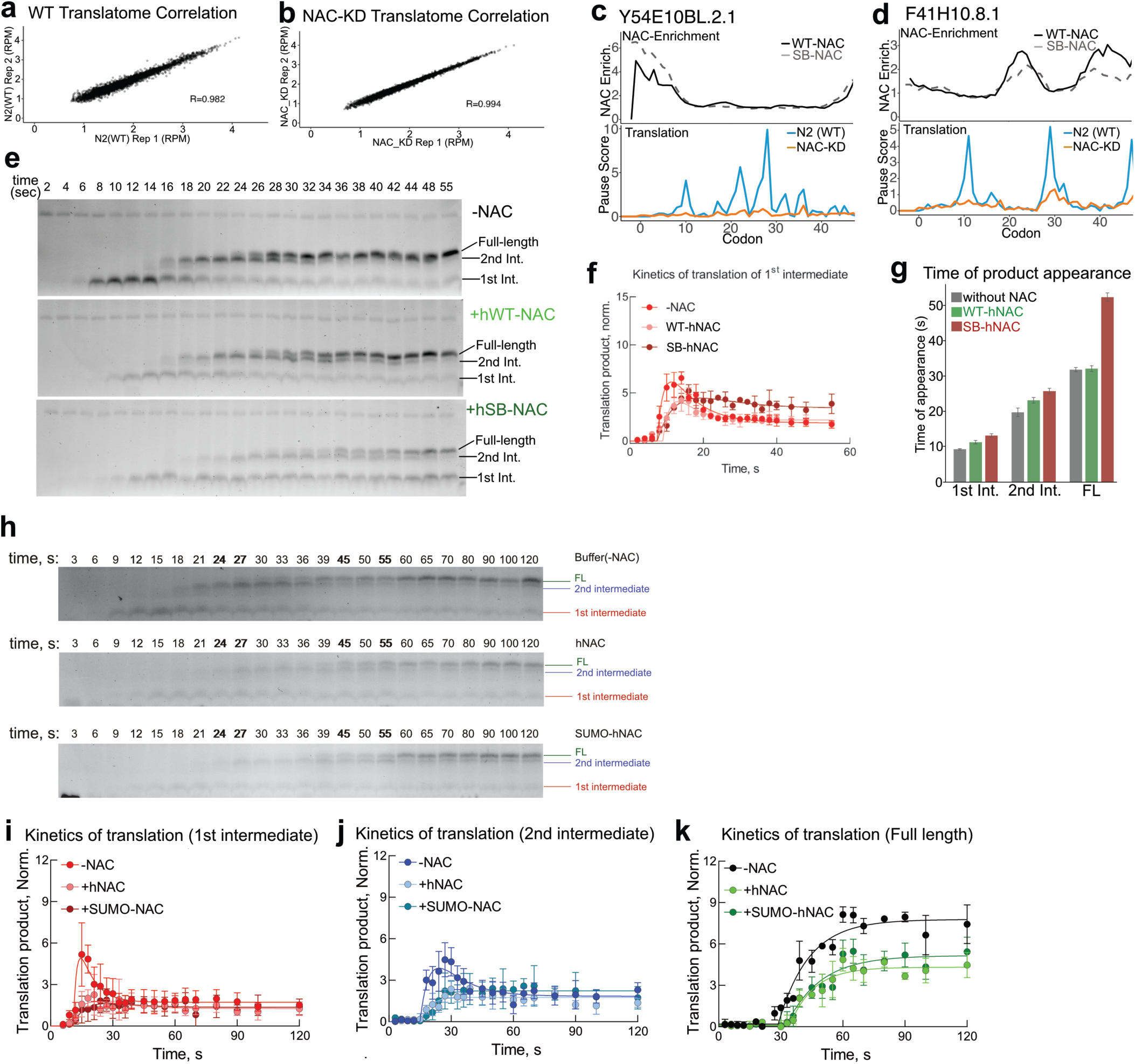
NAC-dependent changes in elongation kinetics. **a-b,** Correlation between replicates from ribosome profiling of wildtype (**a**) or with NAC-KD (**b**). **c-d,** Example proteins that interact with NAC in **Early** (**c**, Y54E10BL.2.1) or **Late** (**d**, F41H10.8.1) region showing NAC enrichment (top) and NAC-dependent slowdowns (bottom). **e-g,** Translation in human reconstituted IVT. Appearance of translation products in the absence or presence of hNAC_WT_/hNAC_ΔN1-67_ over time; SDS_PAGE gel (**e**), accumulation and disappearance of the first intermediate (**f**), and times of product appearance (**g**) calculated from the time courses in panel f and Fig. 5. **h-k,** Effect of NAC on early translation kinetics was monitored as described in Fig. 5d but using a NAC variant with SUMO domain fused to N-terminus of NACβ to prevent insertion into the ribosome exit tunnel. Translation products were separated on a gel as shown in (**h**) and different species (1^st^ intermediate (**i**), 2^nd^ intermediate (**j**), and Full-length (**k**)) were quantified in the absence (-NAC) or in the presence of WT human NAC (+hNAC) or SUMO-fused NAC (+SUMO-NAC) .

**Extended Data Figure 8.**
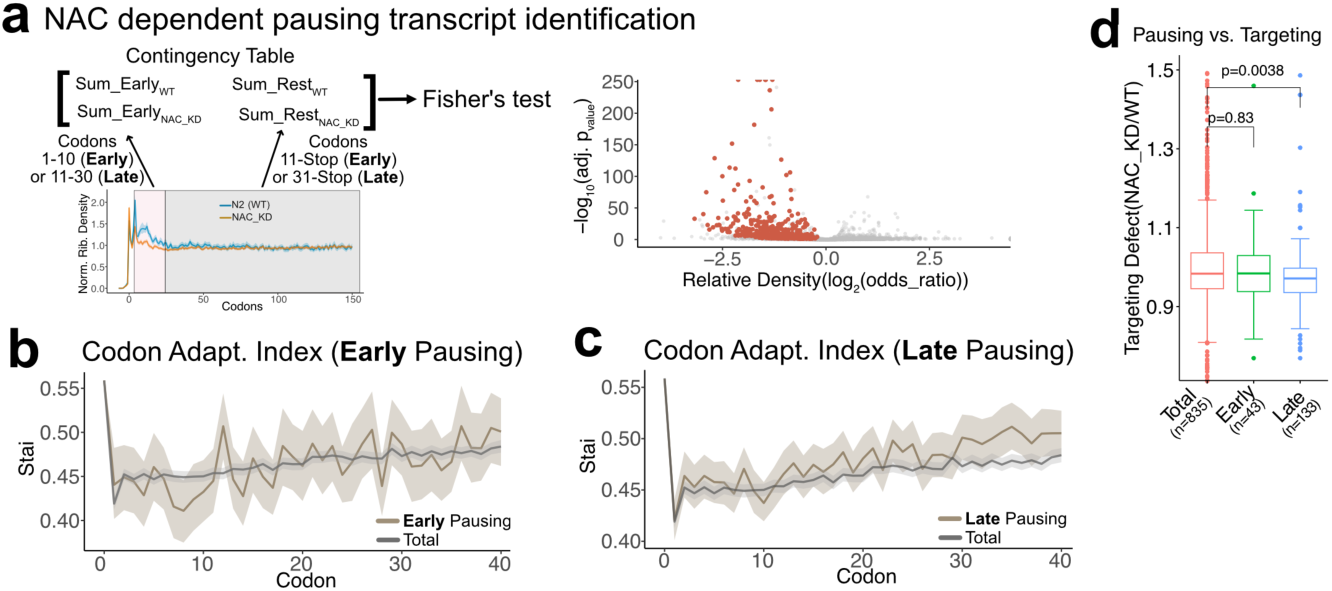
Substrate specific NAC dependent changes in translation speed. **a,** Schematic of identifying transcripts with NAC-dependent changes in local translation. Ribosome density in a specific region was compared to the rest of the transcript between conditions (WT or NAC-KD) to probe transcripts with statistically significant differences using Fisher’s test. **b-c,** Average tRNA adaptation index (stAI)^66^ around start codon for all transcripts (Total, grey), **Early** pausing transcripts (**b**, brown), and **Late** pausing transcripts (**c**, brown). **d,** Targeting defect based on membrane localization of mRNA (Fig. 3o) are compared for all ER/Sec subset that were analyzed (Total), ER/Sec proteins with NAC dependent pausing in ‘Early’ region (Early), or ER/Sec proteins with NAC dependent pausing in ‘Late’ region (Late). Statistical analysis was performed using Fisher’s exact test.

**Extended Data Figure 9.**
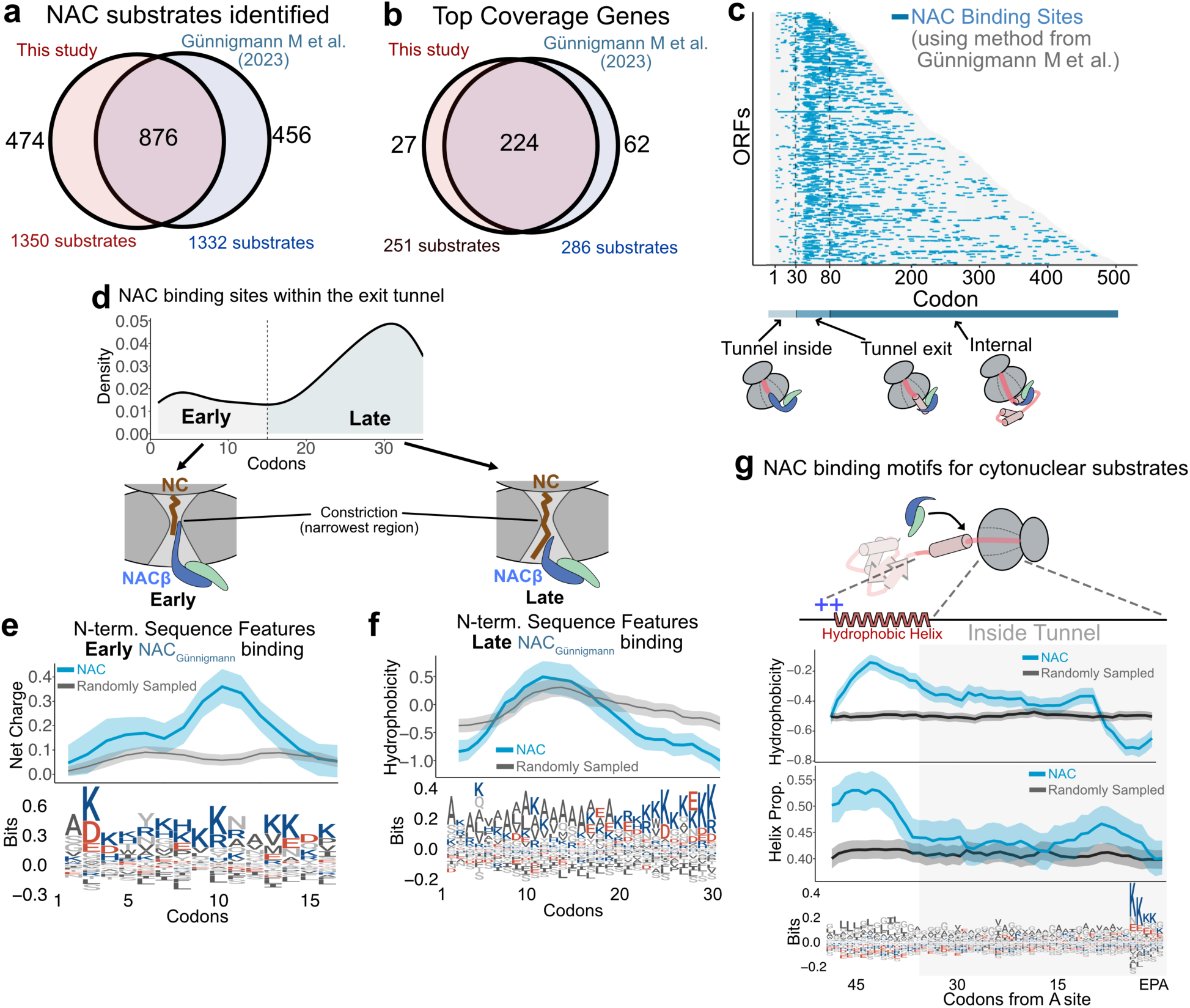
Comparing conclusions from analysis based on transcriptome-wide normalization. Code from^67^ was used to reanalyze the data. **a,** Overlap between NAC substrates identified through two different analysis platforms. **b,** Overlap between subset of genes from **a** that has very good coverage (number of reads). **c,** NAC binding sites mapped to all NAC binding transcripts. **d,** NAC binding site distribution within the first 30 codons of transcripts to show two populations as in the original analysis. **e-f,** Sequence features at the N-terminus of nascent chains with NAC binding sites within the first 15 codons (**e**) or between 15 and 30 codons (**f**). **g,** NAC binding motifs for cytonuclear NAC substrates.

## Notes

### Competing Interest Statement

The authors have declared no competing interest.

## References

1 Mistry, J. et al. Pfam: The protein families database in 2021. Nucleic Acids Res 49, D412–D419 (2021). 10.1093/nar/gkaa913

2 Stein, K. C., Kriel, A. & Frydman, J. Nascent Polypeptide Domain Topology and Elongation Rate Direct the Cotranslational Hierarchy of Hsp70 and TRiC/CCT. Molecular Cell 75, 1117–1130.e1115 (2019). 10.1016/j.molcel.2019.06.036

3 Lewis, T. E. et al. Gene3D: Extensive prediction of globular domains in proteins. Nucleic Acids Res 46, D435–D439 (2018). 10.1093/nar/gkx1069

4 Sillitoe, I. et al. CATH: increased structural coverage of functional space. Nucleic Acids Res 49, D266–D273 (2021). 10.1093/nar/gkaa1079

5 Fukasawa, Y. et al. MitoFates: improved prediction of mitochondrial targeting sequences and their cleavage sites. Mol Cell Proteomics 14, 1113–1126 (2015). 10.1074/mcp.M114.043083

6 Almagro Armenteros, J. J., et al. Detecting sequence signals in targeting peptides using deep learning. Life Sci Alliance 2 (2019). 10.26508/lsa.201900429

7 Jumper, J. et al. Highly accurate protein structure prediction with AlphaFold. Nature 596, 583–589 (2021). 10.1038/s41586-021-03819-2

8 Varadi, M. et al. AlphaFold Protein Structure Database: massively expanding the structural coverage of protein-sequence space with high-accuracy models. Nucleic Acids Res 50, D439–D444 (2022). 10.1093/nar/gkab1061

9 Du, Z. et al. Structure of the human respiratory complex II. Proc Natl Acad Sci U S A 120, e2216713120 (2023). 10.1073/pnas.2216713120

10 Gamerdinger, M. et al. Early Scanning of Nascent Polypeptides inside the Ribosomal Tunnel by NAC. Molecular Cell 75, 996–1006.e1008 (2019). 10.1016/j.molcel.2019.06.030

11 Balchin, D., Hayer-Hartl, M. & Hartl, F. U. In vivo aspects of protein folding and quality control. Science 353, aac4354-aac4354 (2016). 10.1126/science.aac4354

12 Gamerdinger, M. & Deuerling, E. Cotranslational sorting and processing of newly synthesized proteins in eukaryotes. Trends in Biochemical Sciences 49, 105–118 (2024). 10.1016/j.tibs.2023.10.003

13 Morales-Polanco, F., Lee, J. H., Barbosa, N. M. & Frydman, J. Cotranslational Mechanisms of Protein Biogenesis and Complex Assembly in Eukaryotes. Annual Review of Biomedical Data Science 5, 67–94 (2022). 10.1146/annurev-biodatasci-121721-095858

14 Becker, A. H., Oh, E., Weissman, J. S., Kramer, G. & Bukau, B. Selective ribosome profiling as a tool for studying the interaction of chaperones and targeting factors with nascent polypeptide chains and ribosomes. Nature Protocols 8, 2212–2239 (2013). 10.1038/nprot.2013.133

15 Chartron, J. W., Hunt, K. C. L. & Frydman, J. Cotranslational signal-independent SRP preloading during membrane targeting. Nature 536, 224–228 (2016). 10.1038/nature19309

16 Pechmann, S., Willmund, F. & Frydman, J. The ribosome as a hub for protein quality control. Mol Cell 49, 411–421 (2013). 10.1016/j.molcel.2013.01.020

17 Hipp, M. S., Kasturi, P. & Hartl, F. U. The proteostasis network and its decline in ageing. Nature Reviews Molecular Cell Biology 20, 421–435 (2019). 10.1038/s41580-019-0101-y

18 Samatova, E., Komar, A. A. & Rodnina, M. V. How the ribosome shapes cotranslational protein folding. Curr Opin Struct Biol 84, 102740 (2024). 10.1016/j.sbi.2023.102740

19 Stein, K. C. & Frydman, J. The stop-and-go traffic regulating protein biogenesis: How translation kinetics controls proteostasis. Journal of Biological Chemistry 294, 2076–2084 (2019). 10.1074/jbc.REV118.002814

20 Reimann, B. et al. Initial characterization of the nascent polypeptide-associated complex in yeast. Yeast 15, 397–407 (1999). 10.1002/(SICI)1097-0061(19990330)15:5<397::AID-YEA384>3.0.CO;2-U

21 Wiedmann, B. & Prehn, S. The nascent polypeptide-associated complex (NAC) of yeast functions in the targeting process of ribosomes to the ER membrane. FEBS letters 458, 51–54 (1999).

22 Hsieh, H. H., Lee, J. H., Chandrasekar, S. & Shan, S. O. A ribosome-associated chaperone enables substrate triage in a cotranslational protein targeting complex. Nat Commun 11, 5840 (2020). 10.1038/s41467-020-19548-5

23 Jomaa, A. et al. Mechanism of signal sequence handover from NAC to SRP on ribosomes during ER-protein targeting. Science 375, 839–844 (2022). 10.1126/science.abl6459

24 Zhang, Y. et al. NAC functions as a modulator of SRP during the early steps of protein targeting to the endoplasmic reticulum. Molecular Biology of the Cell 23, 3027–3040 (2012). 10.1091/mbc.E12-02-0112

25 Gamerdinger, M., Hanebuth, M. A., Frickey, T. & Deuerling, E. The principle of antagonism ensures protein targeting specificity at the endoplasmic reticulum. Science 348, 201–207 (2015). 10.1126/science.aaa5335

26 Nyathi, Y. & Pool, M. R. Analysis of the interplay of protein biogenesis factors at the ribosome exit site reveals new role for NAC. Journal of Cell Biology 210, 287–301 (2015). 10.1083/jcb.201410086

27 Alamo, M. d., et al. Defining the Specificity of Cotranslationally Acting Chaperones by Systematic Analysis of mRNAs Associated with Ribosome-Nascent Chain Complexes. PLoS Biology 9, e1001100–e1001100 (2011). 10.1371/journal.pbio.1001100

28 Gamerdinger, M. et al. NAC controls cotranslational N-terminal methionine excision in eukaryotes. Science 380, 1238–1243 (2023). 10.1126/science.adg3297

29 Lentzsch, A. M. et al. NAC guides a ribosomal multienzyme complex for nascent protein processing. Nature (2024). 10.1038/s41586-024-07846-7

30 Gamerdinger, M. et al. Mechanism of cotranslational protein N-myristoylation in human cells. Mol Cell 85, 2749–2758 e2748 (2025). 10.1016/j.molcel.2025.06.015

31 Tuller, T. et al. An evolutionarily conserved mechanism for controlling the efficiency of protein translation. Cell 141, 344–354 (2010). 10.1016/j.cell.2010.03.031

32 Verma, M. et al. A short translational ramp determines the efficiency of protein synthesis. Nat Commun 10, 5774 (2019). 10.1038/s41467-019-13810-1

33 Han, Y. et al. Ribosome profiling reveals sequence-independent post-initiation pausing as a signature of translation. Cell Res 24, 842–851 (2014). 10.1038/cr.2014.74

34 Oh, E. et al. Selective ribosome profiling reveals the cotranslational chaperone action of trigger factor in vivo. Cell 147, 1295–1308 (2011). 10.1016/j.cell.2011.10.044

35 Schibich, D. et al. Global profiling of SRP interaction with nascent polypeptides. Nature 536, 219–223 (2016). 10.1038/nature19070

36 Döring, K. et al. Profiling Ssb-Nascent Chain Interactions Reveals Principles of Hsp70-Assisted Folding. Cell 170, 298–311.e220 (2017). 10.1016/j.cell.2017.06.038

37 Menichelli, E., Isel, C., Oubridge, C. & Nagai, K. Protein-induced Conformational Changes of RNA During the Assembly of Human Signal Recognition Particle. Journal of Molecular Biology 367, 187–203 (2007). 10.1016/j.jmb.2006.12.056

38 Fünfschilling, U. & Rospert, S. Nascent polypeptide-associated complex stimulates protein import into yeast mitochondria. Molecular Biology of the Cell 10, 3289–3299 (1999). 10.1091/mbc.10.10.3289

39 George, R., Walsh, P., Beddoe, T. & Lithgow, T. The nascent polypeptide-associated complex (NAC) promotes interaction of ribosomes with the mitochondrial surface in vivo. FEBS Letters 516, 213–216 (2002). 10.1016/S0014-5793(02)02528-0

40 Marc, P. et al. Genome-wide analysis of mRNAs targeted to yeast mitochondria. EMBO reports 3, 159–164 (2002).

41 Lesnik, C., Cohen, Y., Atir-Lande, A., Schuldiner, M. & Arava, Y. OM14 is a mitochondrial receptor for cytosolic ribosomes that supports co-translational import into mitochondria. Nature Communications 5, 5711–5711 (2014).

42 Ponce-Rojas, J. C. et al. αβ′-NAC cooperates with Sam37 to mediate early stages of mitochondrial protein import. The FEBS Journal 284, 814–830 (2017). 10.1111/febs.14024

43 Avendaño-Monsalve, M. C. et al. Positively charged amino acids at the N terminus of select mitochondrial proteins mediate early recognition by import proteins αβ0-NAC and Sam37. Journal of Biological Chemistry 298, 101984–101984 (2022). 10.1016/j.jbc.2022.101984

44 Muthukumar, G. et al. Triaging of alpha-helical proteins to the mitochondrial outer membrane by distinct chaperone machinery based on substrate topology. Mol Cell 84, 1101–1119 e1109 (2024). 10.1016/j.molcel.2024.01.028

45 Eaglesfield, R. & Tokatlidis, K. Targeting and Insertion of Membrane Proteins in Mitochondria. Front Cell Dev Biol 9, 803205 (2021). 10.3389/fcell.2021.803205

46 Akopian, D., Shen, K., Zhang, X. & Shan, S.-o. Signal Recognition Particle: An Essential Protein-Targeting Machine. Annual Review of Biochemistry 82, 693–721 (2013). 10.1146/annurev-biochem-072711-164732

47 Nilsson, I. et al. The Code for Directing Proteins for Translocation across ER Membrane: SRP Cotranslationally Recognizes Specific Features of a Signal Sequence. Journal of Molecular Biology 427, 1191–1201 (2015). 10.1016/j.jmb.2014.06.014

48 Wilson, D. N. & Beckmann, R. The ribosomal tunnel as a functional environment for nascent polypeptide folding and translational stalling. Curr Opin Struct Biol 21, 274–282 (2011). 10.1016/j.sbi.2011.01.007

49 Dao Duc, K., Batra, S. S., Bhattacharya, N., Cate, J. H. D. & Song, Y. S. Differences in the path to exit the ribosome across the three domains of life. Nucleic Acids Res 47, 4198–4210 (2019). 10.1093/nar/gkz106

50 Ito, K. & Chiba, S. Arrest peptides: cis-acting modulators of translation. Annu Rev Biochem 82, 171–202 (2013). 10.1146/annurev-biochem-080211-105026

51 Requiao, R. D., Barros, G. C., Domitrovic, T. & Palhano, F. L. Influence of nascent polypeptide positive charges on translation dynamics. Biochem J 477, 2921–2934 (2020). 10.1042/BCJ20200303

52 McGlincy, N. J. & Ingolia, N. T. Transcriptome-wide measurement of translation by ribosome profiling. Methods 126, 112–129 (2017). 10.1016/j.ymeth.2017.05.028

53 Schuller, A. P., Wu, C. C. C., Dever, T. E., Buskirk, A. R. & Green, R. eIF5A Functions Globally in Translation Elongation and Termination. Molecular Cell 66, 194–205.e195 (2017). 10.1016/j.molcel.2017.03.003

54 Stein, K. C., Morales-Polanco, F., van der Lienden, J., Rainbolt, T. K. & Frydman, J. Ageing exacerbates ribosome pausing to disrupt cotranslational proteostasis. Nature 601, 637–642 (2022). 10.1038/s41586-021-04295-4

55 Yi, S. H. et al. Conformational rearrangements upon start codon recognition in human 48S translation initiation complex. Nucleic Acids Res 50, 5282–5298 (2022). 10.1093/nar/gkac283

56 Gerashchenko, M. V., Peterfi, Z., Yim, S. H. & Gladyshev, V. N. Translation elongation rate varies among organs and decreases with age. Nucleic Acids Res 49, e9 (2021). 10.1093/nar/gkaa1103

57 Agirrezabala, X. et al. A switch from alpha-helical to beta-strand conformation during co-translational protein folding. EMBO J 41, e109175 (2022). 10.15252/embj.2021109175

58 Galmozzi, C. V. et al. Proteome-wide determinants of co-translational chaperone binding in bacteria. Nat Commun 16, 4361 (2025). 10.1038/s41467-025-59067-9

59 Shen, K. et al. Dual Role of Ribosome-Binding Domain of NAC as a Potent Suppressor of Protein Aggregation and Aging-Related Proteinopathies. Molecular Cell 74, 729–741.e727 (2019). 10.1016/j.molcel.2019.03.012

60 Kirstein-Miles, J., Scior, A., Deuerling, E. & Morimoto, R. I. The nascent polypeptide-associated complex is a key regulator of proteostasis. EMBO Journal 32, 1451–1468 (2013). 10.1038/emboj.2013.87

61 Schroeder, A. M. et al. Nascent polypeptide-Associated Complex and Signal Recognition Particle have cardiac-specific roles in heart development and remodeling. PLoS Genetics 18, 1–28 (2022). 10.1371/journal.pgen.1010448

62 Walther, D. M. et al. Widespread Proteome Remodeling and Aggregation in Aging C. elegans. Cell 161, 919–932 (2015). 10.1016/j.cell.2015.03.032

63 Di Fraia, D., et al. Impaired biogenesis of basic proteins impacts multiple hallmarks of the aging brain. bioRxiv (2024). 10.1101/2023.07.20.549210

64 Muslinkina, L. et al. Two independent routes of post-translational chemistry in fluorescent protein FusionRed. Int J Biol Macromol 155, 551–559 (2020). 10.1016/j.ijbiomac.2020.03.244

65 Jaremko, L., Jaremko, M., Giller, K., Becker, S. & Zweckstetter, M. Structure of the mitochondrial translocator protein in complex with a diagnostic ligand. Science 343, 1363–1366 (2014). 10.1126/science.1248725

66 Sabi, R., Volvovitch Daniel, R. & Tuller, T. stAIcalc: tRNA adaptation index calculator based on species-specific weights. Bioinformatics 33, 589–591 (2017). 10.1093/bioinformatics/btw647

67 Gunnigmann, M., Koubek, J., Kramer, G. & Bukau, B. Selective ribosome profiling as a tool to study interactions of translating ribosomes in mammalian cells. Methods Enzymol 684, 1–38 (2023). 10.1016/bs.mie.2022.09.006

68 Brenner, S. THE GENETICS OF CAENORHABDITIS ELEGANS. Genetics 77, 71–94 (1974). 10.1093/genetics/77.1.71

69 Mello, C. & Fire, A. in Methods in Cell Biology Vol. 48 (eds Henry F. Epstein & Diane C. Shakes) 451–482 (Academic Press, 1995).

70 Shemiakina, I. I. et al. A monomeric red fluorescent protein with low cytotoxicity. Nature Communications 3, 1204 (2012). 10.1038/ncomms2208

71 Ketting, R. F., Tijsterman, M. & Plasterk, R. H. Introduction of Double-Stranded RNA in C. elegans by Feeding. Cold Spring Harbor Protocols 2006, pdb. prot4317 (2006).

72 Frokjaer-Jensen, C. et al. Random and targeted transgene insertion in Caenorhabditis elegans using a modified Mos1 transposon. Nat Methods 11, 529–534 (2014). 10.1038/nmeth.2889

73 Schindelin, J., et al. Fiji: an open-source platform for biological-image analysis. Nature Methods 9, 676–682 (2012). 10.1038/nmeth.2019

74 Sharma, A., Mariappan, M., Appathurai, S. & Hegde, R. S. 339–363 (2010).

75 Pestova, T. V. & Hellen, C. U. Reconstitution of eukaryotic translation elongation in vitro following initiation by internal ribosomal entry. Methods 36, 261–269 (2005). 10.1016/j.ymeth.2005.04.004

76 Pisarev, A. V., Unbehaun, A., Hellen, C. U. & Pestova, T. V. Assembly and analysis of eukaryotic translation initiation complexes. Methods Enzymol 430, 147–177 (2007). 10.1016/S0076-6879(07)30007-4

77 Korniy, N. et al. Modulation of HIV-1 Gag/Gag-Pol frameshifting by tRNA abundance. Nucleic Acids Res 47, 5210–5222 (2019). 10.1093/nar/gkz202

78 Park, J. H. et al. Production of active recombinant eIF5A: reconstitution in E.coli of eukaryotic hypusine modification of eIF5A by its coexpression with modifying enzymes. Protein Eng Des Sel 24, 301–309 (2011). 10.1093/protein/gzq110

79 Milon, P. et al. Transient kinetics, fluorescence, and FRET in studies of initiation of translation in bacteria. Methods Enzymol 430, 1–30 (2007). 10.1016/S0076-6879(07)30001-3

80 Kumar, P., Hellen, C. U. & Pestova, T. V. Toward the mechanism of eIF4F-mediated ribosomal attachment to mammalian capped mRNAs. Genes Dev 30, 1573–1588 (2016). 10.1101/gad.282418.116

81 Schagger, H. & von Jagow, G. Tricine-sodium dodecyl sulfate-polyacrylamide gel electrophoresis for the separation of proteins in the range from 1 to 100 kDa. Anal Biochem 166, 368–379 (1987). 10.1016/0003-2697(87)90587-2

82 Schagger, H. Tricine-SDS-PAGE. Nat Protoc 1, 16–22 (2006). 10.1038/nprot.2006.4

83 Osorio, D., Rondón-Villarreal, P. & Torres, R. Peptides: A Package for Data Mining of Antimicrobial Peptides. The R Journal 7 (2015). 10.32614/rj-2015-001

84 Thomsen, M. C. & Nielsen, M. Seq2Logo: a method for construction and visualization of amino acid binding motifs and sequence profiles including sequence weighting, pseudo counts and two-sided representation of amino acid enrichment and depletion. Nucleic Acids Res 40, W281–287 (2012). 10.1093/nar/gks469

85 Dobin, A. et al. STAR: ultrafast universal RNA-seq aligner. Bioinformatics 29, 15–21 (2013). 10.1093/bioinformatics/bts635

86 McGuffin, L. J., Bryson, K. & Jones, D. T. The PSIPRED protein structure prediction server. Bioinformatics 16, 404–405 (2000). 10.1093/bioinformatics/16.4.404

87 Goldberg, T. et al. LocTree3 prediction of localization. Nucleic Acids Res 42, W350–355 (2014). 10.1093/nar/gku396

